# Direct RNA nanopore sequencing of full-length coronavirus genomes provides novel insights into structural variants and enables modification analysis

**DOI:** 10.1101/483693

**Authors:** Adrian Viehweger, Sebastian Krautwurst, Kevin Lamkiewicz, Ramakanth Madhugiri, John Ziebuhr, Martin Hölzer, Manja Marz

## Abstract

Sequence analyses of RNA virus genomes remain challenging due to the exceptional genetic plasticity of these viruses. Because of high mutation and recombination rates, genome replication by viral RNA-dependent RNA polymerases leads to populations of closely related viruses, so-called ‘quasispecies’. Standard (short-read) sequencing technologies are ill-suited to reconstruct large numbers of full-length haplotypes of (i) RNA virus genomes and (ii) subgenome-length (sg) RNAs comprised of noncontiguous genome regions. Here, we used a full-length, direct RNA sequencing (DRS) approach based on nanopores to characterize viral RNAs produced in cells infected with a human coronavirus.

Using DRS, we were able to map the longest (∼26 kb) contiguous read to the viral reference genome. By combining Illumina and nanopore sequencing, we reconstructed a highly accurate consensus sequence of the human coronavirus (HCoV) 229E genome (27.3 kb). Furthermore, using long reads that did not require an assembly step, we were able to identify, in infected cells, diverse and novel HCoV-229E sg RNAs that remain to be characterized. Also, the DRS approach, which circumvents reverse transcription and amplification of RNA, allowed us to detect methylation sites in viral RNAs. Our work paves the way for haplotype-based analyses of viral quasispecies by demonstrating the feasibility of intra-sample haplotype separation.

Even though several technical challenges remain to be addressed to exploit the potential of the nanopore technology fully, our work illustrates that direct RNA sequencing may significantly advance genomic studies of complex virus populations, including predictions on long-range interactions in individual full-length viral RNA haplotypes.

## Background

Coronaviruses (subfamily *Coronavirinae*, family *Coronaviridae*, order *Nidovirales*) are enveloped positive-sense (+) single-stranded (ss) RNA viruses that infect a variety of mammalian and avian hosts and are of significant medical and economic importance, as illustrated by recent zoonotic transmissions from diverse animal hosts to humans^1,2^. The genome sizes of coronaviruses (∼30 kb) exceed those of most other RNA viruses. Coronaviruses use a special mechanism called discountinuous extension of minus strands^3,4^ to produce a nested set of 5’- and 3’-coterminal subgenomic (sg) mRNAs that carry a common 5’ leader sequence that is identical to the 5’ end of the viral genome^5,6^. These sg mRNAs contain a different number of open reading frames (ORFs) that encode the viral structural proteins and several accessory proteins. With very few exceptions, only the 5’-located ORF (which is absent from the next smaller sg mRNA) is translated into protein (Fig. 1).

**Figure 1:**
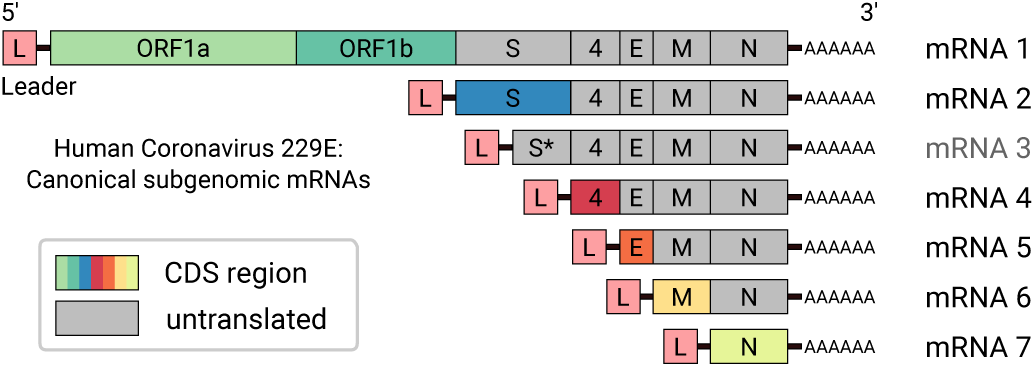
Scheme of genomic and subgenomic (sg) RNAs produced in HCoV-229E-infected cells^7,8^. Translation of the 5’-terminal ORF(s) of the respective RNA gives rise to the various viral structural and nonstructural proteins (indicated by different colors). RNA 3 is considered defective and unlikely to be translated into protein. Each RNA has a 3’ poly(A) tail and carries a 5’-leader sequence that is identical to the 5’-end of the genome. In a process called discontinuous extension of negative strands, negative-strand RNA synthesis is attenuated at specific genomic positions. After transfer of the nascent strand to an upstream position on the template, RNA synthesis is continued by copying the 5’-leader sequence. As a result, the 3’-ends of coronavirus minus-strand RNAs are equipped with the complement of the 5’-leader. The latter process is guided by base-pairing interactions between the transcription-regulating sequence (TRS) located immediately downstream of the leader sequence (TRS-L) at the 5’-end of the genome and the sequence complement of a TRS located upstream of one of the ORFs present in the 3’-proximal genome region (TRS-B).

In HCoV-229E-infected cells, at total of 7 major viral RNAs are produced. The viral genome (RNA 1) is occasionally referred to as mRNA 1 because it (also) has an mRNA function. In its 5’-terminal region, the genome RNA contains two large ORFs, 1a and 1b, that encode the viral replicase polyproteins 1a and 1ab. mRNAs 2, 4, 5, 6, and 7 are used to produce the S protein, accessory protein 4, E protein, M protein and N protein, respectively. The 5’-ORF present in RNA 3 starts contains the central and 3’ regions of the S gene. Although this sg RNA has been consistently identified in HCoV-229E-infected cells, its mRNA function has been disputed and there is currently no evidence that this RNA is translated into protein^7–9^.

Like many other +RNA viruses, coronaviruses show high rates of recombination^10–12^. In fact, the mechanism to produce 5’ leader-containing sg mRNAs represents a prime example for copy-choice RNA recombination that, in this particular case, is guided by complex RNA-RNA interactions involving the transcription-regulating sequence (TRS) core sequences and likely requires additional interactions of viral proteins with specific RNA signals. In other virus systems, RNA recombination has been shown to generate ‘transcriptional units’ that control the expression of individual components of the genome^13^. The mechanisms involved in viral RNA recombination are diverse and may even extend to non-replicating systems^14^. In the vast majority of cases, recombination results in defective RNA (dRNA) copies that lack essential cis-active elements and thus cannot be replicated. In other cases, functional recombinant RNA with new properties, such as the ability to replicate in a new host, may emerge^15–18^. In yet other cases, defective interfering RNAs (DI-RNAs) may be produced. These defective (subgenome-length) RNAs contain all the cis-acting elements required for efficient replication by a helper virus polymerase and, therefore, represent parasitic RNAs that compete for components of the viral replication/transcription complex with non-defective viral RNAs^19^.

To elucidate the many facets of recombination and to determine full-length haplotypes of, for example, virus mutants/variants in complex viral populations (quasispecies), long-read sequencing has become the method of choice. Short-read second-generation sequencing technologies – such as IonTorrent and Illumina – are restricted by read length (200-400 nucleotides^20^). For example, the use of highly fragmented viral RNAs considerably complicates the investigation of haplotypes^21,22^. Since the nested coronavirus mRNAs are almost identical to the original genome sequence, short-read data can usually not be unambiguously assigned to specific sg RNA species.

In this study, we performed direct RNA sequencing (DRS) on an array of nanopores, as developed by Oxford Nanopore Technologies (ONT)^23^. Nanopore sequencing does not have a limited reading length but is limited only by fragmentation of the input material^24–26^. Further, by using DRS, we avoid several drawbacks of previous sequencing methods, in particular cDNA synthesis and amplification of the input material. Thus, for example, cDNA synthesis can create artificial RNA-RNA chimerae^27^ that are difficult to discriminate from naturally occurring chimerae (such as spliced RNAs). Also, amplification prior to sequencing would remove all RNA modifications from the input material, whereas the nanopore sequencing technology preserves these modifications^23,28^.

Recently, nanopore sequencing has been used for metagenomic forays into the virosphere^29^ and studies focusing on transmission routes^30,31^. Furthermore, viral transcriptomes have been investigated using nanopore sequencing of cDNA^32–35^, being subject to bias from reverse transcription and amplification. Other studies used DRS to study the human poly(A) transcriptome^36^ and the transcriptome of DNA viruses such as HSV^37^. Furthermore, the genome of Influenza A virus has been completely sequenced in its original form using direct RNA sequencing^38^.

In the present study, we sequenced one of the largest known RNA genomes, that of HCoV-229E, a member of the genus Alphacoronavirus, with a genome size of about 27,300 nt, in order to assess the complex architectural details for viral sg RNAs produced in cells infected with recombinant HCoV-229E. Using DRS, we aim to capture complete viral mRNAs, including the full coronavirus genome, in single contiguous reads. Sequence analysis of thousands of full-length sg RNAs will allow us to determine the architectures (including leader-body junction sites) of the major viral mRNAs. In addition, this approach provides insight into the diversity of additional HCoV-229E sg RNAs, probably including DI-RNAs. Further, we aim to assess whether RNA modifications can be called directly from the raw nanopore signal of viral molecules without prior *in vitro* treatment, as has been shown for DNA^39,40^.

## Results

### Full genome sequencing without amplification

We sequenced total RNA samples obtained from Huh-7 cells infected with serially passaged recombinant human coronaviruses: wild-type (WT) HCoV-229E, HCoV-229E_SL2-SARS-CoV, and HCoV-229E_SL2-BCoV, respectively. In the latter two viruses, a conserved stem-loop structure (SL2) residing in the HCoV-229E 5’-UTR was replaced with the equivalent SL2 element from SARS-CoV and BCoV, respectively^41^. Total RNA samples obtained for the latter two (chimeric) viruses were pooled prior to sequence analysis. Hereafter, we refer to the first sample as WT RNA and to the second (pooled) sample as SL2 RNA (see Methods and Materials).

We performed two direct RNA sequencing runs (one per sample) on a MinION nanopore sequencer. As shown in Table 1, we achieved a throughput of 0.237 and 0.282 gigabases with 225 k and 181k reads for the WT and SL2 sample, respectively. See Supplemental Fig. 1 A for an overview of the read length distribution. For the WT and SL2 samples, 33.2% and 35.9% of the reads mapped to the reference HCoV-229E sequences, respectively. 15.8% and 10.2% respectively mapped to the yeast enolase 2 mRNA sequence, a calibration strand added during the library preparation, while 47.4% and 52.7% could be attributed to human host cell RNA. `minimap2` did not align the remaining 3.50% and 1.11% of reads. Using BLAST^42^ against the `nt` database, 18.1% and 20.7% of these reads can be attributed again to HCoV, human or yeast. As reads which were not aligned by `minimap2` were mostly very short (median <= 200 nt), of poor basecalling quality and represented only 0.62% and 0.15% of total nucleotides respectively, we decided to only use the higher quality reads that minimap2 could align. (see Supplemental Fig. 2 for detailed statistics). The visualized raw voltage signal of a nanopore read is commonly called ‘squiggle’ (see Supplemental Fig. 3). Different from all previous sequencing technologies, nanopore sequencing preserves the information about base modifications in the raw signal^23^. However, one of the biggest challenges is the accurate mapping of the raw voltage signal to bases (’base calling’).

**Table 1:**
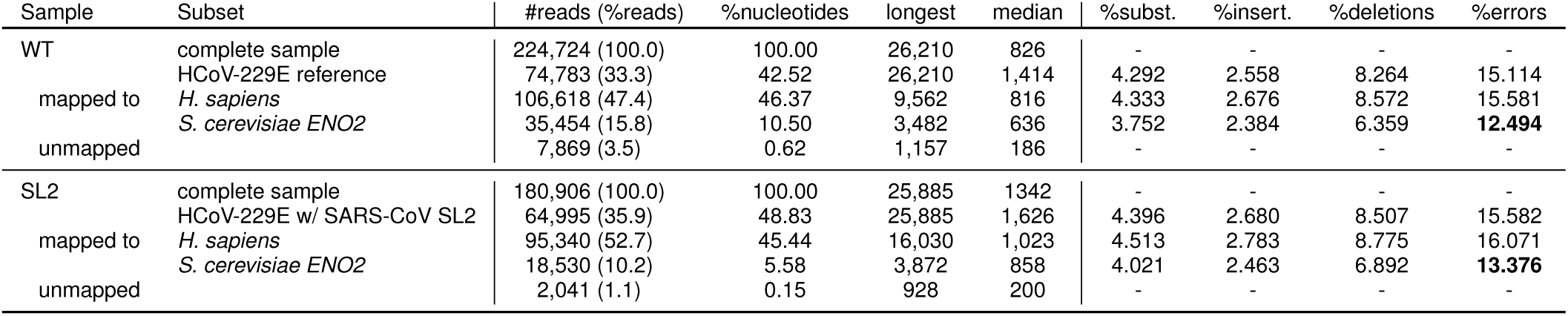
Sequencing and error statistics. Both samples mainly contain HCoV-229E reads, but also host (*Homo sapiens*) transcripts and *S. cerevisiae* enolase 2 (*ENO2*) mRNA (which was used as a calibration standard added during library preparation). Half of the sequencing errors were deletion errors, probably resulting to a large extent from basecalling at homopolymer stretches. The *S. cerevisiae* enolase 2 mRNA reads display an overall reduced error rate because the `Albacore` basecaller was trained on this calibration strand. Note that all error rates report differences to the reference genome and thus include actual genetic variation.

As expected for nanopore DRS^23,38^, reads had a median uncorrected error rate of about 15% for human and virus reads, while basecalling errors were reduced for yeast *ENO2* mRNA reads, as the basecaller was trained on this calibration strand (see Table 1). This included gaps but omitted dis-continuous sites longer than six nucleotides since they indicated recombination. Half of all errors were deletions. In addition, we found that more than half of all single nucleotide deletions occur in homopolymers, and most of these streches that coincide with a deletion are three or more nucleotides long (see Supplemental Fig. 4). A quarter of the errors were substitutions, which we argue are largely due to modified bases that impede the basecaller’s ability to assign bases correctly.

The HCoV-229E genome was 99.86% covered, with a large coverage bias towards both ends (see Fig. 1 and 2). The high coverage of the 3’-end reflects the higher abundance of mRNAs produced from the 3’-terminal genome regions and is a result of the discontinuous transcription mechanism employed by coronaviruses and several other nidoviruses^5,43,44^. The 3’-coverage is further increased by the directional sequencing that starts from the mRNA 3’-terminal poly(A) tail. Also, the observed coverage bias for the very 5’-end results from the coronavirus-specific transcription mechanism because all viral mRNAs are equipped with the 65-nt 5’-leader sequence derived from the 5’-end of the genome. The remainder of the high 5’-coverage bias likely reflects the presence of high numbers of DI-RNAs in which 5’- and 3’-proximal genomic sequences were fused, probably resulting from illegitimate recombination events as shown previously for other coronaviruses^10,12,45^. For the WT and SL2 samples, 38.37% and 16.32% were split-mapped, respectively. Of these, only 278 and 181 had multiple splits. The considerably larger fraction of split reads in the WT sample is explained by the high abundance of potential DI-RNA molecules, see Fig. 2 (c).

**Figure 2:**
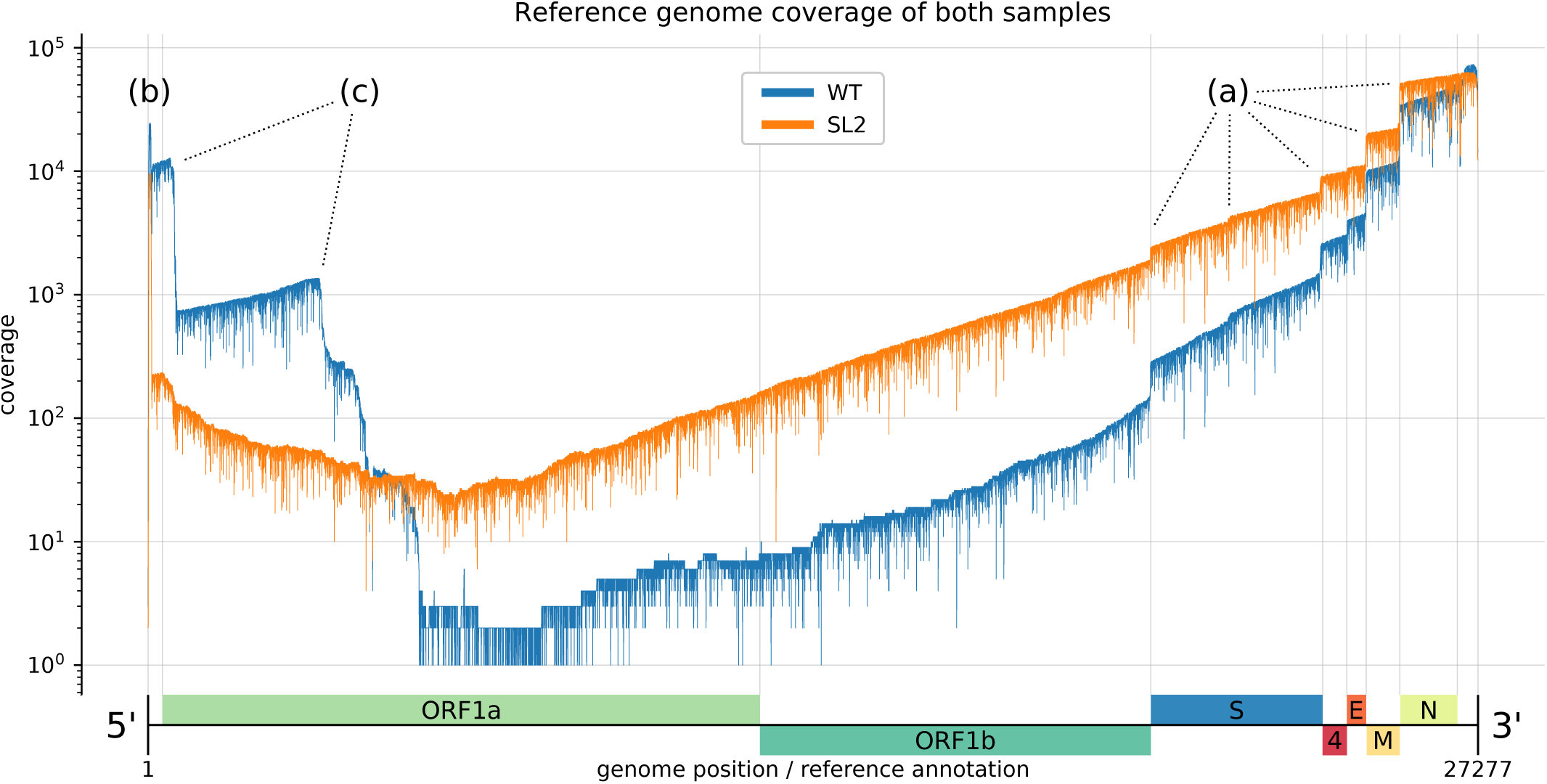
Reference genome coverage of the HCoV-229E WT sample (blue) and the SL2 sample (orange) based on alignments with `minimap2`. There is an inverse correlation between sg RNA abundance and length. **(a)** Notable vertical ‘steps’ in the coverage correspond to borders expected for the canonical sg RNAs (see Fig. 1). **(b)** The presence of the leader sequence (ca. 65 nt) in canonical sg RNAs gives rise to the sharp coverage peak at the 5’-end. **(c)** We also observed unexpected ‘steps’, especially in the WT sample (blue). We hypothesize that the sequences correspond to DI-RNA molecules that may arise by recombination at TRS-like sequence motifs as well as other sites displaying sequence similarities that are sufficient to support illegitimate recombination events (see Fig. 3). We attribute the difference in the observed (non-canonical) recombination sites between the two samples to biological factors that we either did not control for or do not know (see also legend to Fig. 3).

An alignment of the longest reads from both samples to the HCoV-229E reference indicates that they represent near complete virus genomes (Supplemental Fig. 1 B). The observed peaks in the aligned reads length distribution (see Fig. 4) corresponded very well with the abundances of the known mRNAs produced in HCoV-229E-infected cells^7–9^ (see Fig. 1). Alignment of the reads to these canonical mRNA sequences confirmed these observed abundances (Supplemental Fig. 5).

**Figure 3:**
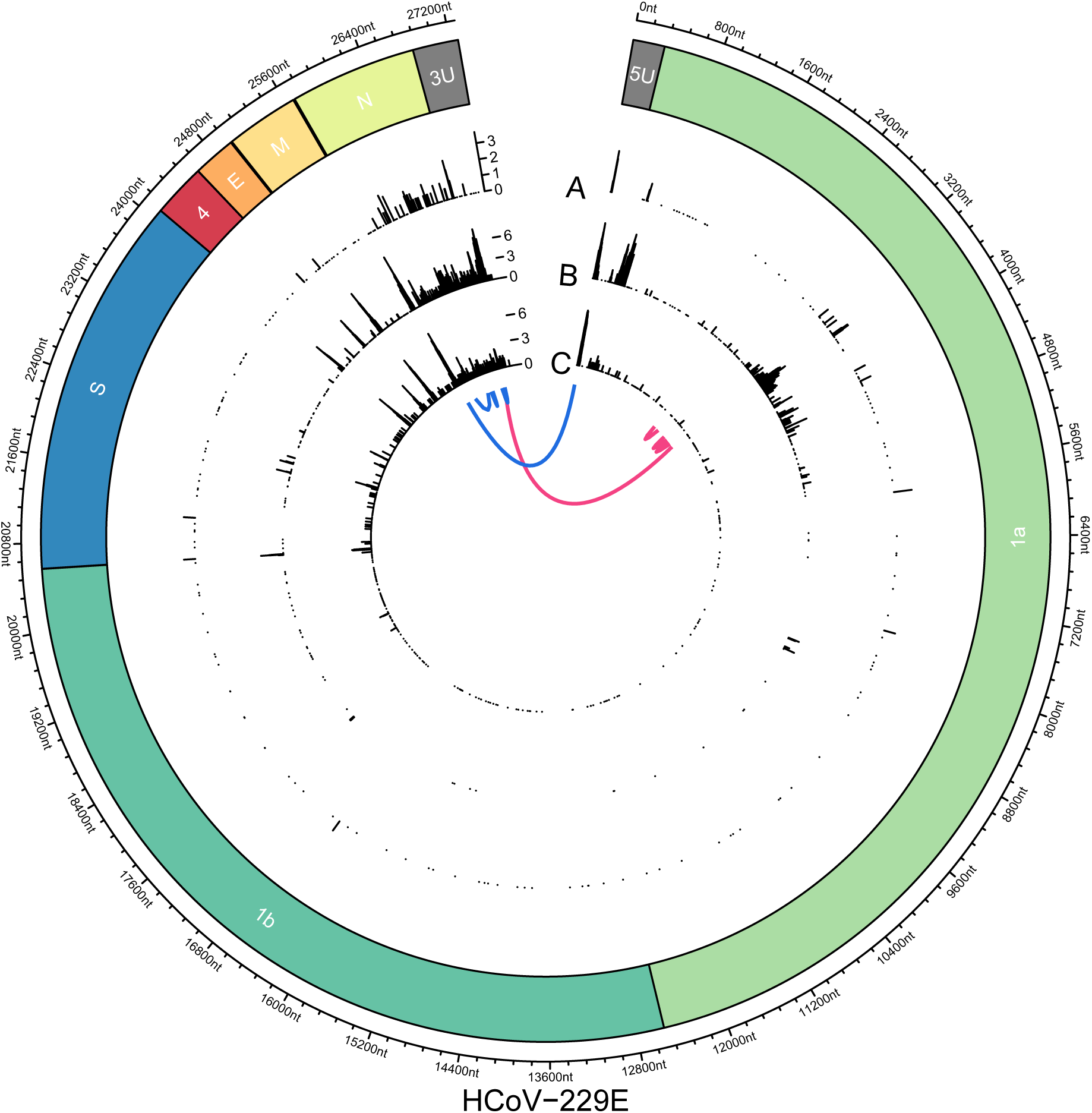
Joining of non-contiguous genome sequences in sg RNAs identified in HCoV-229E-infected cells. On the **circular axis** the annotations of the reference genome (including 5’-UTR (5U) and 3’-UTR (3U)) are shown. Genomic positions of ‘discontinuous sites’ identified in Illumina reads **(A, outer track)**, nanopore reads of sample HCoV-229E WT **(B, middle track)** and nanopore reads of SL2 sample **(C, inner track)** reveal multiple recombination sites across the whole genome. An aggregation of recombination sites can be observed in the region that encodes the viral N protein. Furthermore, clear recombination sites can be seen at intergenic boundaries and at the 5’- and 3’-UTRs, with the former corresponding to the boundary between the leader sequence and the rest of the genome. Another prominent cluster can be observed in ORF1a in the WT nanopore sample, but not in SL2. This cluster is supported by the WT Illumina data, excluding sequencing bias as a potential source of error. We hypothesize that since samples WT and SL2 were obtained from non-plaque-purified serially passaged virus populations derived from *in vitro*-transcribed genome RNAs transfected into Huh-7 cells, differences in the proportion of full-length transcripts versus abortive transcripts could translate into different patterns of recombination. Generally, nanopore-based sequencing allows more detailed analysis of recombination events due to the long read length. Even complex isoforms such as two exemplary reads, each with four discontinuous segments, can be observed (blue and pink).

**Figure 4:**
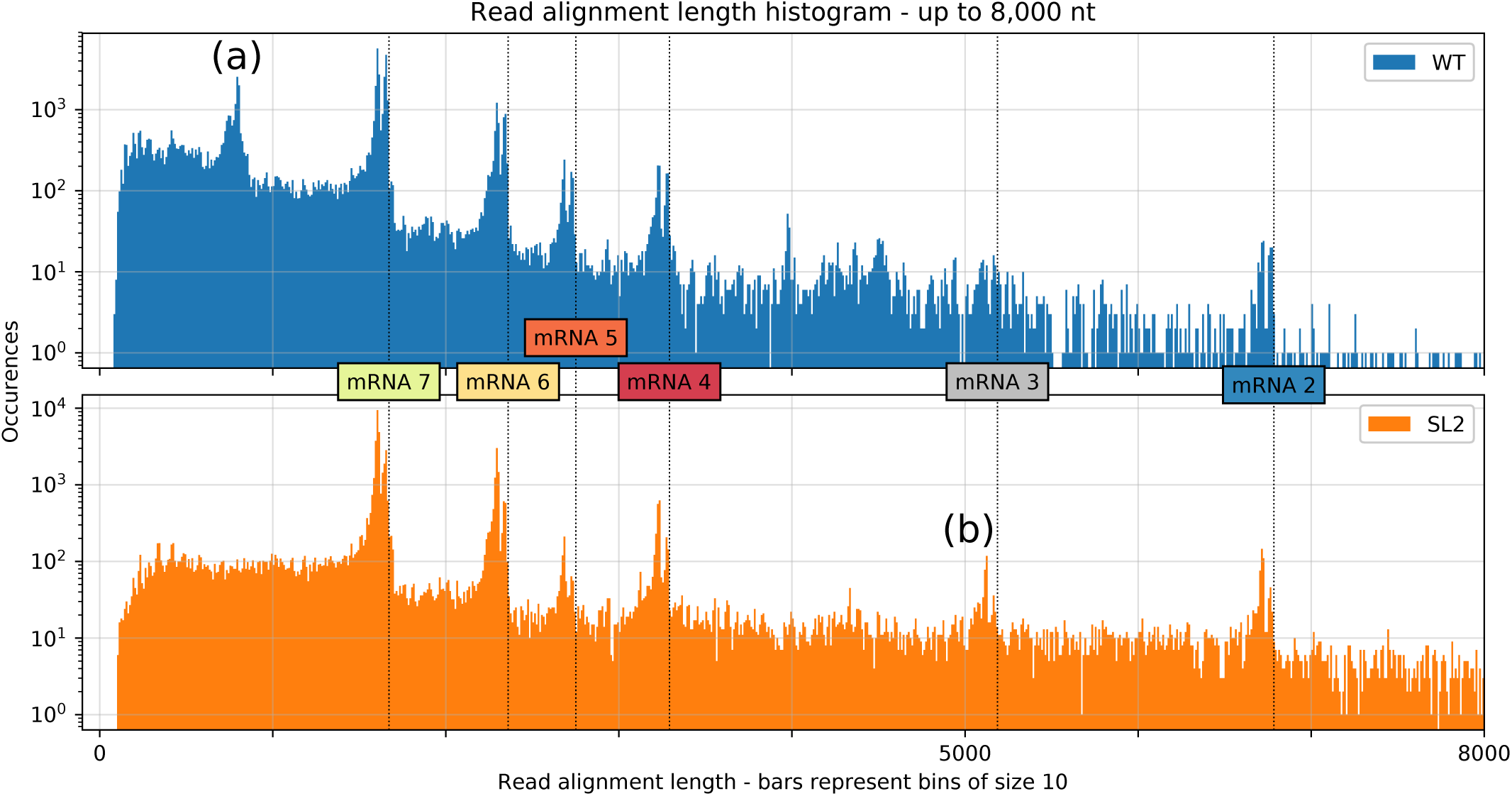
Distribution of aligned read lengths up to 8,000 nt for both samples based on alignments with `minimap2`. We observed clusters that correspond well with the transcript lengths expected for canonical mRNAs (vertical dotted lines). Alignment of the reads to these canonical mRNA sequences confirmed these observed abundances (Supplemental Fig. 5). The distribution shows double peaks at the cluster positions because reads corresponding to mRNAs often miss the leader sequence, possibly due to basecalling or mapping errors. We also observed additional clusters that likely correspond to highly abundant DI-RNAs. **(a)** indicates a cluster in the WT sample that respresents chimeric ∼820-nt sg RNAs comprised of both 5’- and 3’-terminal genome regions (∼540 nt and ∼280 nt, respectively). We propose the template switch for this transcript occurs around position 27,000 and RNA synthesis resumes at around position 540 (see corresponding peaks in Fig. 3 track B). **(b)** indicates a cluster from the SL2 sample that contains transcripts with an approximate length of 5,150 nt which represents mRNA 3 (see Fig. 1). These transcripts are probably formed due to a transcription stop at a TRS motif around position 22,150 (see corresponding peak in Fig. 3 track C).

The median read length for the combined set of reads from both samples was 826 nt, with a maximum of 26,210 nt, covering 99.86% of the 27,276-nt-long virus genome, missing only 21 nt at the 5’-end, 15 nt at the 3’-end and those nucleotides that correspond to the skewed error distribution, with 5.7 percentage points more deletions than insertions (see Tab. 1). The median read length might sound short, however most of the viral RNAs (including many DI-RNAs) identified in HCoV-229E-infected cells were below 2,000 nt in length. Furthermore, this number nearly doubles the longest read length that can be obtained with short-read sequencing methods. We observed an abundance of very short reads, representing the 3’ (poly-(A)) end of the genome. This could be an artifact of RNA degradation, although we cannot estimate the exact fraction of affected transcripts. Because sequencing starts at the poly-(A) tail, fragmented RNA will not be sequenced beyond any 3’ break point. It is thus best to minimize handling time during RNA extraction and library preparation. Innovations in these fields will directly translate into larger median read lengths.

We obtained 99.15% and 98.79% identity in both samples (WT, SL2) respectively with the help of the consensus caller `Ococo`^46^ using the reference genomes and all reads mapping to it. We attempted a standard long-read assembly using `Canu`^47^, which yielded unusable results (WT: 389 contigs, longest 13 kb, all other <4 kb; SL2: 517 contigs, all <6 kb). We think that current nanopore-only assembly tools are not equipped to handle special read data sets such as those originating from a small RNA virus genome. In addition, we assembled WT and SL2 consensus sequences using Nanopore and Illumina data with `HG-CoLoR`^48^ in an approach that uses long nanopore reads to traverse an assembly graph constructed from short Illumina reads. We thereby recovered 99.57% of the reference genome in a single contiguous sequence at 99.90% sequence identity to reference using this approach with the single longest read from the SL2 sample. This hybrid approach illustrates how short- and long-read technologies can be combined to reconstruct long transcripts accurately, which will greatly facilitate studies of haplotypes.

### Uncharacterized subgenomic and defective interfering RNAs

In addition to the leader-to-body junctions expected for the canonical sg mRNAs 2, 4, 5, 6, and 7, we observed a high number of recombination sites (Fig. 3) which were consistently found in our samples but have not been described previously (Fig. 3). In this study, we defined a recombination site as any site that flanks more than 100 consecutive gaps, as determined in a discontinuous mapping (’spliced’ mapping). While there is currently no consensus on how to define such sites, we believe this to be a conservative definition, as this type of pattern is unlikely to result from e.g. miscalled homopolymer runs which, in our experience, typically affect less than 10 consecutive bases. We observe all known canonical HCoV-229E mRNAs at their expected lengths, including the (presumably) non-coding mRNA 3 (Fig. 4).

The aligned reads distribution revealed clusters for all known mRNAs which closely fit the expected molecule lengths (Fig. 4). The cluster positions show double peaks with a consistent distance of ∼65 nt, i.e. the length of the leader sequence. We observed that the 5’-end of reads has larger-than-average error rates and is often missing nucleotides (see Supplemental Fig. 8 for detailed statistics). This might be due to a bias of the basecaller towards the end of reads. This is plausible, because the underlying classification algorithm is a bidirectional (i.e. forward and backward looking) long-short-term memory neural network (LSTM). The mapping algorithm was often unable to align these erroneous 5’-ends, leading to soft-clipped bases. Thus, for many reads representing canonical mRNAs, the included leader sequence was not aligned, which gives rise to the secondary peak at each cluster position. We also observed additional clusters which likely correspond to highly abundant dRNAs (Fig. 4).

We also observed several unexpected recombination sites, e.g. at positions 3,000 to 4,000 (within ORF1a, see Fig. 3). These sites were confirmed by both nanopore and Illumina sequencing. They had a high read support and defined margins, suggesting a specific synthesis/amplification of these sg RNAs which, most likely, represent DI-RNAs. Since DI-RNAs are byproducts of viral replication and transcription, they present a larger diversity than the canonical viral mRNAs^49–53^.

Nanopore sequencing captures recombination events far better than Illumina, which allowed us to identify even complex sg RNAs (composed of sequences derived from more than 2 non-contiguous genome regions) at much higher resolution: For example, we found sg RNAs with up to four recombination sites in the 5’- and 3’-terminal genome regions (Fig. 3).

### Consistent 5mC methylation signatures of coronavirus RNA

Nanopore sequencing preserves information on nucleotide modifications. Using a trained model, DNA and RNA modifications such as 5mC methylation could be identified (Fig. 5). To assess the false positive rate (FPR) of the methylation calling, we used an unmethylated RNA calibration standard (RCS) as a negative control which was added in the standard library preparation protocol for DRS. We considered a position to be methylated if at least 90% of the reads showed a methylation signal for this particular position. Using this threshold, the estimated FPR was calculated to be below 5%. Our experimental setup did not include a positive methylation control.

**Figure 5:**
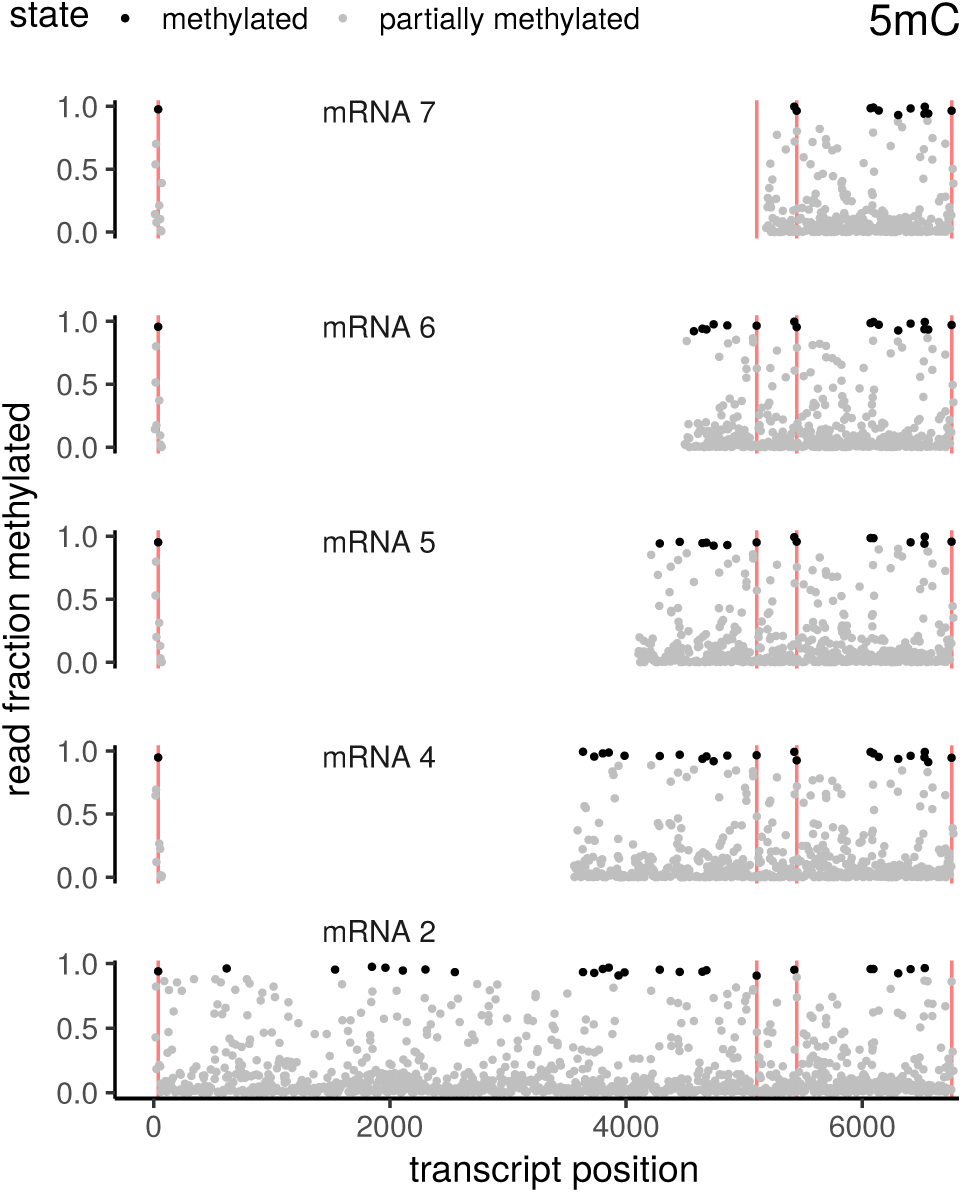
5mC methylation of various annotated coronavirus mRNAs. The transcripts have been aligned such that corresponding genomic positions can be found in the same vertical column across facets. Both the leader sequence (including the TRS) as well as the nested subgenomic sequences show consistent patterns of methylation across transcripts. Exemplary positions that display consistent methylation across all investigated transcripts are marked as red vertical lines. Note that while coronavirus recombination uses two TRS, the resulting transcript has only one TRS, because of self-similarity based pairing of the TRS. Positions have been labelled ‘methylated’ if at least 90% of the reads show a methylation signal. Using this threshold, the estimated FPR is below 5%.

When analyzing 5mC methylation across various RNAs, we observed consistent patterns (Figure 5) that were reproducible for the corresponding genomic positions in different RNAs, suggesting that the methylation of coronavirus RNAs is sequence-specific and/or controlled by RNA structural elements. Methylated nucleotides could be identified across the genome, both in the leader sequence and in the body regions of viral mRNAs.

While the overall methylation pattern looks similar between subgenomic RNAs and the negative control (see Supplemental Fig. 9), we nevertheless find consistent methylation across different subgenomic RNA “types", i.e. methylated positions of mRNA 2 are mirrored in mRNA 4 etc.

## Discussion

We identified highly diverse sg RNAs in coronavirus-infected cells, with many sg RNAs not corresponding to the known canonical mRNAs. These ‘non-canonical’ sg RNAs had abundant read support and full-length sequences could be obtained for most of these RNAs.

As indicated above, only 12% of the sg RNAs were found to conform to our current understanding of discontinuous mRNA transcription in coronaviruses, resulting in mRNAs that (all) carry an identical 5’-leader sequence that is fused to the 3’-coding (’body’) sequence of the respective mRNA. We however believe that 12% represents an underestimate because a large number of sg RNAs were probably omitted from the analysis: (1) RNA molecules degrade rapidly under laboratory conditions, even when handled carefully. The resulting fragments will only be sequenced if they contain a poly(A) tail. (2) The high sequencing error may introduce mismappings, especially for low-quality reads. These reads would not be assigned to the canonical model under our assumptions because of the high number of mismatches. However, we think the associated bias is low, because `minimap2` is very robust against high error rates and because the reads are very long, thus ensuring that the mapper has sufficient aggregate information on a given read to position it very reliably on a reference. (3) The library preparation protocol for DRS includes the ligation of adapters via a T4 ligase. Any ligase could potentially introduce artificial chimera, although we did not investigate this systematically. Again, we think that this does not affect our results substantially: First, this bias is random and it seems unlikely that we would observe the very same RNA ‘isoform’ many times if it was created by random ligation. Second, many ‘isoforms’ that we observed only once (e.g. those colored pink and blue in Figure 3) were structured plausibly: They contained a leader sequence and had recombined at expected (self-similar) sites corresponding to putative or validated TRSs, with downstream sequences being arranged in a linear 5’-3’-order. (4) Finally, it is important to note that the RNA used for DRS was isolated from cells infected with a serially passaged pool of recombinant viruses rescued after transfection of *in vitro*-transcribed genome-length (27.3 kb) RNAs. Transfection of preparations of *in vitro*-transcribed RNA of this large size likely included a significant proportion of abortive transcripts that lacked varying parts of the 3’ genome regions, rendering them dysfunctional. It is reasonable to suggest that the presence of replication-incompetent RNAs lacking essential 3’-terminal genome regions may have triggered recombination events resulting in the emergence of DI-RNAs that contained all the 5’- and 3’-terminal cis-active elements required for RNA replication, but lacked non-essential internal genome regions. Upon serial passaging of the cell culture supernants for 21 times, DI-RNAs may have been enriched, especially in the HCoV-229E (wild-type) sample (Figure 3). Comparative DRS analyses of RNA obtained from cells infected with (i) plaque-purified HCoV-229E and (ii) newly rescued recombinant HCoV-229E (without prior plaque purification), respectively, would help to address the possible role of prematurely terminated *in vitro* transcripts produced from full-length cDNA in triggering the large number of DI-RNAs observed in our study.

Although, for the above reasons, the low percentage of canonical mRNAs (12%) in our samples likely represents an underestimate, our study may stimulate additional studies, for example to revisit the production of mRNAs from non-canonical templates^54,55^. Also, it is worth mentioning in this context that, for several other nidoviruses, such as murine hepatitis virus (MHV), bovine coronavirus (BCoV), and arteriviruses, evidence has been obtained that sg RNA transcription may also involve non-canonical TRS motifs ^56–60^.

The majority of sg RNAs (other than mRNAs) we found in our samples likely represent DI-RNAs, which are a common occurrence in coronavirus *in vitro* studies^19^.

To our knowledge, this study is the first to perform RNA modification calling without prior treatment of the input sample. It only relies on the raw nanopore signal. While DNA modifications such as 5mC methylation have been explored extensively^61^, less is known about RNA modifications^62^, the importance of which is debated^63^. We found consistent 5mC methylation patterns across viral RNAs when tested at a FPR below 5%. We were not able to assess the sensitivity and specificity of the methylation calling due to the absence of a positive control group, which was beyond the scope of this study. RNA is known to have many different modifications, and we expect the presence of these on Coronavirus sgRNAs^64^ too. However, to our knowledge no comprehensive data exists on prior expectations for such modifications in Coronaviruses, which might or might not correspond to those observed in e.g. humans.

In addition, we observed that the software used^39^ will likely present high error rates in regions of low coverage or where the underlying reference assembly is erroneous. This is because the resquiggle algorithm – upon which this method is based – has to align the raw nanopore read signal to the basecalled read sequence (see Supplemental Fig. 6). This is necessary to test the raw signal against learned modification models, of which at the time of manuscript preparation (May 2019) only 5mC was implemented for RNA. Nevertheless, new options to call these modifications at an acceptable error rate without any RNA pretreatment is a powerful method.

The validity of the methylation signal should be confirmed in future studies using e.g. bisulfite sequencing. Ideally, this validation should start from *in vitro*, synthetic transcripts where modified bases have been inserted in known positions. Furthermore, RNA modification detection from single-molecule sequencing is a current bioinformatic frontier, and algorithms and tools are under active development. We showed that consistent 5mC methylation patterns were seen across different subgenomic RNAs. However, the overall pattern of the methylation calls between subgenomic RNAs and the negative control was very similar. At a false positive rate of 5%, the RNA modifications we identified are supported by their consistent occurence. However, we cannot rule out that instead, the observed pattern might be caused by an alignment artifact. In the employed methylation calling algorithm, the raw signal is aligned to the nucleotide sequence after basecalling. If there is a systematic bias in this alignment, and certain sequence motifs cause a consistent mapping mismatch, this mismatch could lead to false positive methylated sites. This is, because in these positions the signal would deviate from the expected one due to the misalignment, and not due to methylation. In future experiments this can be decided using a positive control in the form of an RNA transcript with known 5mC methylated sites. However, even if we are in its early stages, the reading of RNA modifications from the read signal has great potential to elucidate viral biology.

We were able to reconstruct accurate consensus sequences, both for the Illumina and nanopore data. We also demonstrated that individual transcripts can be characterized. More problematic was the resolution of quasispecies in our experimental setup. Although DRS allowed us to confirm the presence of each of the two heterologous SL2 structures present in the SL2 sample, this was only possible for subgenome-length (DI-) RNAs. It appears that the high error rate of more than 10% was a critical limitation when analyzing the SL2 region located at the extreme 5’-terminal end of the 27.3 kb genome RNA. This high error rate made variant calling difficult, particularly under low-coverage conditions, as was the case in our analyses of the 5’-UTR of genome-length RNA (results not included). The current generation of long-read assemblers is not well suited to reconstruct many viral genome architectures, such as nested ones. The development of specialized assemblers would be of great help in virology projects.

We used a hybrid error correction method (`HG-CoLoR`^48^) that uses Illumina data to correct read-level errors. However, it remains questionable whether the corrected read sequence is truly representative of the ground truth read sequence. Signal-based correction methods such as `Nanopolish`^65^ may be more promising, however, at the time of manuscript writing (May 2019) correction on direct RNA data has not been implemented. We expect this to become available in the near future. Combined with the ever-increasing accuracy of the nanopore technology, we think this method might be able to study quasispecies soon.

There are recombination events observed in the Illumina data that were not detected in the Nanopore data. These are likely caused by misalignment of the short single-end reads (50 nt). A minimum of only 10 nt was required for mapping on either end of the gapped alignment. This was a trade-off between sensitivity to identify recombination sites and unspecific mapping.

In this work, we demonstrated the potential of long-read data as produced by nanopore sequencing. We were able to directly sequence the RNA molecules of two different samples of one of the largest RNA virus genomes known to date. We showed how very large RNA genomes and a diverse set of sg RNAs with complex structures can be investigated at high resolution without the need for a prior assembly step and without the bias introduced by cDNA synthesis that is typically required for transcriptome studies.

The detail and quality of the available data still require significant bioinformatic expertise as the available tooling is still at an early development stage. However, the technological potential of nanopore sequencing for new insights into different aspects of viral replication and evolution is very promising.

Future studies should investigate both strands of the coronavirus transcriptome. Studies focusing on RNA modifications need to employ well-defined positive and negative controls to assess the error rate of the current software alternatives. Also, the DRS method will be extremely powerful if it comes to analyzing the nature and dynamics of specific haplotypes in coronavirus populations under specific selection pressures, for example mutations and/or drugs affecting replication efficiency and others.

Our work also serves as a proof-of-concept demonstrating that consistent RNA modifications can be detected using nanopore DRS.

To fully exploit the potential of DRS, several improvements are needed: First and foremost, a significant reduction of the currently very high per-read error rate is crucial. This is especially problematic in studies focusing on intra-sample heterogeneity and haplotypes. Secondly, protocols that limit RNA degradation during library preparation would be of great value. This could be achieved by shortening the library protocol. To limit the cost of DRS, barcoded adapters would be desirable. On the bioinformatics side, the basecaller for DRS data is still at an early stage and, for example, cannot accurately call the poly(A) regions as well as the RNA-DNA-hybrid adapter sequences. Further basecalling errors likely result from RNA modifications, which need to be modelled more accurately. However, once these limitations will be fixed, the use of nanopore-based DRS can be expected to greatly advance our understanding of the genomics of virus populations and their multiple haplotypes.

## Methods

### RNA virus samples

The two total RNA samples used in this study for DRS (ONT MinION) and Illumina sequencing were prepared at 24 h post infection from Huh-7 cells infected at an MOI of 3 with recombinant HCoV-229E WT, HCoV-229E_SL2-SARS-CoV and HCoV-229E_SL2-BCoV, respectively^41^. Prior to sequence analysis, the two RNA samples obtained from HCoV-229E_SL2-SARS-CoV- and HCoV-229E_SL2-BCoV-infected cells were pooled (SL2 sample, see Supplemental Fig. 7).

Generation of recombinant viruses and total RNA isolation were carried out as described previously^41^. Briefly, full-length cDNA copies of the genomes of HCoV-229E (GenBank accession number NC_002645), HCoV-229E_SL2-SARS-CoV and HCoV-229E_SL2-BCoV, respectively, were engineered into recombinant vaccinia viruses using previously described methods^66–68^. Next, full-length genomic RNAs of HCoV-229E, HCoV-229E_SL2-SARS-CoV and HCoV-229E_SL2-BCoV, respectively, were transcribed in vitro using purified ClaI-digested genomic DNA of the corresponding recombinant vaccinia virus as a template. 1.5 µg of full-length viral genome RNA, along with 0.75 µg of in vitro-transcribed HCoV-229E nucleocapsid protein mRNA, were used to transfect 1 *×* 10^6^ Huh-7 cells using the TransIT^®^ mRNA transfection kit according to the manufacturer’s instructions (Mirus Bio LLC). At 72 h post transfection (p.t.), cell culture supernatants were collected and serially passaged in Huh-7 cells for 21 (WT) or 12 times (HCoV-229E_SL2-SARS-CoV and HCoV-229E_SL2-BCoV), respectively.

### Nanopore sequencing and long-read assessment

For nanopore sequencing, 1 µg of RNA in 9 µl was carried into the library preparation with the Oxford nanopore direct RNA sequencing protocol (SQK-RNA001). All steps were followed according to the manufacturer’s specifications. The library was then loaded on an R9.4 flow cell and sequenced on a MinION device (Oxford Nanopore Technologies). The sequencing run was terminated after 48 h.

The raw signal data was basecalled using `Albacore` (v2.2.7, available through the Oxford Nanopore community forum). Due to the size of the raw signal files, only the basecalled data were deposited at the Open Science Framework (OSF; doi.org/10.17605/OSF.IO/UP7B4).

While it is customary to remove adapters after DNA sequencing experiments, we did not perform this pre-processing step. The reason is that the sequenced RNA is attached to the adapter molecule via a DNA linker, effectively creating a DNA-RNA chimera. The current basecaller – being trained on RNA – is not able to reliably translate the DNA part of the sequence into base space, which makes adapter trimming based on sequence distance unreliable. However, we found that the subsequent mapping is very robust against these adapter sequences. All mappings were performed with `minimap2`^69^ (v2.8-r672) using the ‘spliced’ preset without observing the canonical `GU…AG` splicing motif (parameter `-u n`), and *k*-mer size set to 14 (`-k 14`).

Raw reads coverage and sequence identity to the HCoV-229E reference genome (WT: GenBank, NC_002645.1; SL2: stem loop 2 sequence replaced with SARS-CoV SL2 sequence) were determined from mappings to the references produced by `minimap2`. Read origin and sequencing error statistics were assessed by mapping the reads simultaneously with `minimap2` to a concatenated mock-genome consisting of HCoV-229E (WT and SL2 variants respectively), yeast enolase 2 mRNA (calibration strand, GenBank, NP_012044.1), and the human genome (GRCh38). Identity and error rates are the number of matching nucleotides (or number of nucleotide substitutions, insertions or deletions) divided by the total length of the alignment including gaps from indels.

Consensus calling of nanopore reads was performed with `Ococo`^46^ (v0.1.2.6). The minimum required base quality was set to 0 in order to avoid gaps in low coverage domains.

We used the hybrid error correction tool `HG-CoLoR`^48^ in conjunction with the Illumina HiSeq short-read data sets of both samples to reduce errors in all reads that exceed 20k nt in length. The program builds a *de Bruijn* graph from the near noise-free short-read data and then substitutes fragments of the noisy long reads with paths found in the graph which correspond to that same fragment of the sequence. `HG-CoLoR` was run with default parameters except for the maximum order of the *de Bruijn* graph, which was set to 50 in order to fit the length of the short reads.

### Illumina HiSeq sequencing and assembly

Illumina short-read sequencing was performed using the TruSeq RNA v2 kit to obtain RNA from species with poly(A) tails and without any strand information. The three samples (WT, SL2_SARS-CoV, SL2_BCoV) selected for this study were prepared on a HiSeq 2500 lane and sequenced with 51 cycles. After demultiplexing, 23.2, 22.0, and 23.8 million single-end reads were obtained for the WT and the two SL2 samples, respectively. The raw sequencing data was deposited at the OSF (doi.org/10.17605/OSF.IO/UP7B4).

### Characterization of transcript isoforms and subgenomic RNAs

We first defined TRS as a 8-mer with a maximum Hamming distance of 2 from the motif UCUCAACU. We then searched the HCoV-229E reference genome (GenBank, NC_002645.1) for all matching 8-mers. We then synthesized sg RNAs *in silico* as follows: For each pair of complementary 8-mers (5’-TRS, 3’-TRS) we accepted at most 1 mismatch to simulate base pairing under a stable energy state. We then joined two reference subsequences for each pair or TRS: First, the 5’-end up to but not including the 5’-TRS. Second, the 3’-end of the reference genome including the 3’-TRS and excluding the poly(A) tail.

This way we obtained about 5,000 candidate sg RNAs. To validate them, we mapped the nanopore reads to these ‘mock’ sg RNAs in a non-discontinuous manner, all i.e. reads had to map consecutively without large gaps. To count as a putative hit, 95% of the read length had to uniquely map to a given mock transcript, and the mock transcript could not be more than 5% longer than the read.

We only considered putative hits as plausible if they had a read support of at least 5. With this threshold, we aim to balance the sensitivity of finding plausible novel transcripts with a need to control the number of false positives.

### Identification of 5mC methylation

We used `Tombo` (v1.3, default parameters)^39^ to identify signal level changes corresponding to 5mC methylation (see Supplemental Fig. 6).

To assess the false positive rate (FPR) of the methylation calling, we used an RNA calibration standard (RCS) as a negative control. It is added in the standard library preparation protocol for direct RNA sequencing. This mRNA standard is derived from the yeast enolase II (YHR174W) gene^70^ and is produced using an *in vitro* transcription system. As a consequence, the mRNA standard is not methylated.

For a conservative resquiggle match score of 1.3 (part of the `Tombo` algorithm, default setting for RNA) and a methylation threshold of 0.9, the FPR was 4.67%, which met our requirement that the FPR be smaller than 5%. Our experimental setup did not include a positive methylation control.

## Data access

Both the short-read cDNA (Illumina) and the base-called long-read RNA (ONT) data are available from the European Nucleotide Archive (ENA) under accession PRJEB33797, as well as from the Open Science Framework repository UP7B4. Raw long-read data in fast5 format are also available from this OSF repository. All analysis code has been deposited in the same OSF repository and is also available in Supplemental_Code.zip from the supplemental material.

## Acknowledgements

We sincerely thank Celia Diezel for technical assistance in nanopore sequencing. We thank Ivonne Görlich and Marco Groth from the Core Facility DNA sequencing of the Leibniz Institute on Aging – Fritz Lipmann Institute in Jena for their help with Illumina sequencing. We also thank Nadja Karl (Medical Virology, Giessen) for excellent technical assistence. MH appreciates the support of the Joachim Herz Foundation by the add-on fellowship for interdisciplinary life science.

This work was supported by BMBF – InfectControl 2020 (03ZZ0820A) (KL, MM) and is part of the Collaborative Research Centre AquaDiva (CRC 1076 AquaDiva) of the Friedrich Schiller University Jena, funded by the Deutsche Forschungsgemeinschaft (DFG); supporting MH. The study is further supported by DFG TRR 124 “FungiNet”, INST 275/365-1, B05 (MM). The work of JZ was supported by the DFG (SFB 1021-A01 and KFO 309-P3).

We used `Inkscape` version 0.92.1 (available from inkscape.org) to finalize our figures for publication.

## Authors’ contributions

AV developed the experimental design for sequencing with nanopores. AV, SK and KL analyzed and interpreted the data. JZ, RM, MH and MM were major contributors for discussion and in writing the manuscript. All authors wrote, commented, edited and approved the final manuscript.

## References

[1] Vineet D Menachery, Rachel L Graham, and Ralph S Baric. Jumping species-a mechanism for coronavirus persistence and survival. Curr Opin Virol, 23:1–7, 2017. ISSN 1879-6265. doi: 10.1016/j.coviro.2017.01.002.

[2] Rahul Vijay and Stanley Perlman. Middle east respiratory syndrome and severe acute respiratory syndrome. Curr Opin Virol, 16:70–76, 2016. ISSN 1879-6265. doi: 10.1016/j.coviro.2016.01.011.

[3] S G Sawicki and D L Sawicki. Coronaviruses use discontinuous extension for synthesis of subgenome-length negative strands. Adv Exp Med Biol, 380:499–506, 1995. ISSN 0065-2598.

[4] S G Sawicki and D L Sawicki. A new model for coronavirus transcription. Adv Exp Med Biol, 440:215–219, 1998. ISSN 0065-2598.

[5] Stanley G Sawicki, Dorothea L Sawicki, and Stuart G Siddell. A contemporary view of coronavirus transcription. J Virol, 81:20–29, 2007. ISSN 0022-538X. doi: 10.1128/JVI.01358-06.

[6] Sonia Zuniga, Isabel Sola, Sara Alonso, and Luis Enjuanes. Sequence motifs involved in the regulation of discontinuous coronavirus subgenomic RNA synthesis. J Virol, 78:980–994, 2004. ISSN 0022-538X.

[7] T Raabe, B Schelle-Prinz, and S G Siddell. Nucleotide sequence of the gene encoding the spike glycoprotein of human coronavirus HCV 229E. J Gen Virol, 71 (Pt 5):1065–1073, 1990. ISSN 0022-1317. doi: 10.1099/0022-1317-71-5-1065.

[8] S S Schreiber, T Kamahora, and M M Lai. Sequence analysis of the nucleocapsid protein gene of human coronavirus 229E. Virology, 169:142–151, 1989. ISSN 0042-6822.

[9] Volker Thiel, Konstantin A Ivanov, Akos Putics, Tobias Hertzig, Barbara Schelle, Sonja Bayer, Benedikt Weissbrich, Eric J Snijder, Holger Rabenau, Hans Wilhelm Doerr, Alexander E Gorbalenya, and John Ziebuhr. Mechanisms and enzymes involved in SARS coronavirus genome expression. J Gen Virol, 84:2305–2315, 2003. ISSN 0022-1317. doi: 10.1099/vir.0.19424-0.

[10] T Furuya, T B Macnaughton, N La Monica, and M M Lai. Natural evolution of coronavirus defective-interfering RNA involves RNA recombination. Virology, 194:408–413, 1993. ISSN 0042-6822. doi: 10.1006/viro.1993.1277.

[11] M M Lai. Rna recombination in animal and plant viruses. Microbiol Rev, 56:61–79, 1992. ISSN 0146-0749.

[12] C L Liao and M M Lai. Rna recombination in a coronavirus: recombination between viral genomic RNA and transfected RNA fragments. J Virol, 66:6117–6124, 1992. ISSN 0022-538X.

[13] Edward C Holmes. The evolution and emergence of RNA viruses. Oxford University Press, 2009.

[14] Andreas Gallei, Alexander Pankraz, Heinz-Jürgen Thiel, and Paul Becher. RNA recombination in vivo in the absence of viral replication. J Virol, 78:6271–6281, 2004. ISSN 0022-538X. doi: 10.1128/JVI.78.12.6271-6281.2004.

[15] Ruey-Yi Chang, Martin A Hofmann, Phiroze B Sethna, and David A Brian. A cis-acting function for the coronavirus leader in defective interfering RNA replication. J Virol, 68(12):8223–8231, 1994.

[16] Ruey-Yi Chang, Rajesh Krishnan, and David A Brian. The UCUAAAC promoter motif is not required for high-frequency leader recombination in bovine coronavirus defective interfering RNA. J Virol, 70(5):2720–2729, 1996.

[17] Cary G Brown, Kimberley S Nixon, Savithra D Senanayake, and David A Brian. An RNA stem-loop within the bovine coronavirus nsp1 coding region is a cis-acting element in defective interfering RNA replication. J Virol, 81:7716–7724, July 2007. ISSN 0022-538X. doi: 10.1128/JVI.00549-07.

[18] Kortney M Gustin, Bo-Jhih Guan, Agnieszka Dziduszko, and David A Brian. Bovine coronavirus nonstructural protein 1 (p28) is an RNA binding protein that binds terminal genomic cis-replication elements. J Virol, 83(12):6087–6097, 2009.

[19] Kunj B Pathak and Peter D Nagy. Defective interfering RNAs: foes of viruses and friends of virologists. Viruses, 1(3):895–919, 2009.

[20] Martin Hölzer and Manja Marz. Software Dedicated to Virus Sequence Analysis “Bioinformatics Goes Viral". Adv Virus Res, 2017.

[21] M A Nowak. What is a quasispecies? Trends Ecol Evol, 7: 118–121, 1992. ISSN 0169-5347. doi: 10.1016/0169-5347(92)90145-2.

[22] Jasmijn A Baaijens, Amal Zine El Aabidine, Eric Rivals, and Alexander Schönhuth. *De novo* assembly of viral quasispecies using overlap graphs. Genome Res, 27:835–848, 2017. ISSN 1549-5469. doi: 10.1101/gr.215038.116.

[23] Daniel R Garalde, Elizabeth A Snell, Daniel Jachimowicz, Botond Sipos, Joseph H Lloyd, Mark Bruce, Nadia Pantic, Tigist Admassu, Phillip James, Anthony Warland, Michael Jordan, Jonah Ciccone, Sabrina Serra, Jemma Keenan, Samuel Martin, Luke McNeill, E Jayne Wallace, Lakmal Jayasinghe, Chris Wright, Javier Blasco, Stephen Young, Denise Brocklebank, Sissel Juul, James Clarke, Andrew J Heron, and Daniel J Turner. Highly parallel direct RNA sequencing on an array of nanopores. Nat Methods, 15:201–206, 2018. ISSN 1548-7105. doi: 10.1038/nmeth.4577.

[24] Alexander S Mikheyev and Mandy MY Tin. A first look at the Oxford Nanopore MinION sequencer. Mol Ecol Resour, 14(6): 1097–1102, 2014.

[25] Eng Wee Chua and Pei Yuen Ng. MinION: A Novel Tool for Predicting Drug Hypersensitivity? Front Pharmacol, 7:156, 2016. ISSN 1663-9812. doi: 10.3389/fphar.2016.00156.

[26] Miten Jain, Hugh E Olsen, Benedict Paten, and Mark Akeson. The oxford nanopore minION: delivery of nanopore sequencing to the genomics community. Genome Biol, 17:239, 2016. ISSN 1474-760X. doi: 10.1186/s13059-016-1103-0.

[27] Søren M Karst, Morten S Dueholm, Simon J McIlroy, Rasmus H Kirkegaard, Per H Nielsen, and Mads Albertsen. Retrieval of a million high-quality, full-length microbial 16S and 18S rRNA gene sequences without primer bias. Nat Biotechnol, 36:190–195, 2018. ISSN 1546-1696. doi: 10.1038/nbt.4045.

[28] Andrew M Smith, Miten Jain, Logan Mulroney, Daniel R Garalde, Mark Akeson, Alla Mikheenko, Gleb Valin, Andrey Prjibelski, Vladislav Saveliev, and Alexey Gurevich. Reading canonical and modified nucleotides in 16S ribosomal RNA using nanopore direct RNA sequencing. bioRxiv, 32(21):132274, 2017.

[29] Joanna Warwick-Dugdale, Natalie Solonenko, Karen Moore, Lauren Chittick, Ann C Gregory, Michael J Allen, Matthew B Sullivan, and Ben Temperton. Long-read metagenomics reveals cryptic and abundant marine viruses. bioRxiv, page 345041, 2018.

[30] Joshua Quick, Nicholas J Loman, Sophie Duraffour, Jared T Simpson, Ettore Severi, Lauren Cowley, Joseph Akoi Bore, Raymond Koundouno, Gytis Dudas, Amy Mikhail, Nobila Oué-draogo, Babak Afrough, Amadou Bah, Jonathan Hj Baum, Beate Becker-Ziaja, Jan-Peter Boettcher, Mar Cabeza-Cabrerizo, Alvaro Camino-Sanchez, Lisa L Carter, Juiliane Doerrbecker, Theresa Enkirch, Isabel Graciela García Dorival, Nicole Hetzelt, Julia Hinzmann, Tobias Holm, Liana Eleni Kafetzopoulou, Michel Koropogui, Abigail Kosgey, Eeva Kuisma, Christopher H Logue, Antonio Mazzarelli, Sarah Meisel, Marc Mertens, Janine Michel, Didier Ngabo, Katja Nitzsche, Elisa Pallash, Livia Victoria Patrono, Jasmine Portmann, Johanna Gabriella Repits, Natasha Yasmin Rickett, Andrea Sachse, Katrin Singethan, Inês Vitoriano, Rahel L Yemanaberhan, Elsa G Zekeng, Racine Trina, Alexander Bello, Amadou Alpha Sall, Ousmane Faye, Oumar Faye, N’Faly Magassouba, Cecelia V Williams, Victoria Amburgey, Linda Winona, Emily Davis, Jon Gerlach, Franck Washington, Vanessa Monteil, Marine Jourdain, Marion Bererd, Alimou Camara, Hermann Somlare, Abdoulaye Camara, Marianne Gerard, Guillaume Bado, Bernard Baillet, Déborah Delaune, Koumpingnin Yacouba Nebie, Abdoulaye Diarra, Yacouba Savane, Raymond Bernard Pallawo, Giovanna Jaramillo Gutierrez, Natacha Milhano, Isabelle Roger, Christopher J Williams, Facinet Yattara, Kuiama Lewandowski, Jamie Taylor, Philip Rachwal, Daniel Turner, Georgios Pollakis, Julian A Hiscox, David A Matthews, Matthew K O’Shea, Andrew McD Johnston, Duncan Wilson, Emma Hutley, Erasmus Smit, Antonino Di Caro, Roman Woelfel, Kilian Stoecker, Erna Fleischmann, Martin Gabriel, Simon A Weller, Lamine Koivogui, Boubacar Diallo, Sakoba Keita, Andrew Rambaut, Pierre Formenty, Stephan Gunther, and Miles W Carroll. Real-time, portable genome sequencing for ebola surveillance. Nature, 530:228–232, February 2016. ISSN 1476-4687. doi: 10.1038/nature16996.

[31] Nuno R Faria, Josh Quick, IM Claro, Julien Theze, Jacqueline G de Jesus, Marta Giovanetti, Moritz UG Kraemer, Sarah C Hill, Allison Black, Antonio C da Costa, et al. Establishment and cryptic transmission of Zika virus in Brazil and the Americas. Nature, 546 (7658):406, 2017.

[32] Norbert Moldován, Dóra Tombácz, Attila Szűcs, Zsolt Csabai, Michael Snyder, and Zsolt Boldogkői. Multi-platform sequencing approach reveals a novel transcriptome profile in pseudorabies virus. Front Microbiol, 8:2708, 2018.

[33] Dóra Tombácz, Zsolt Csabai, Attila Szűcs, Zsolt Balázs, Norbert Moldován, Donald Sharon, Michael Snyder, and Zsolt Boldogkői. Long-read isoform sequencing reveals a hidden complexity of the transcriptional landscape of herpes simplex virus type 1. Front Microbiol, 8:1079, 2017. ISSN 1664-302X. doi: 10.3389/fmicb.2017.01079.

[34] Norbert Moldován, Zsolt Balázs, Dóra Tombácz, Zsolt Csabai, Attila Szűcs, Michael Snyder, and Zsolt Boldogkői. Multiplatform analysis reveals a complex transcriptome architecture of a circovirus. Virus Res, 237:37–46, 2017. ISSN 1872-7492. doi: 10.1016/j.virusres.2017.05.010.

[35] Norbert Moldován, Dóra Tombácz, Attila Szűcs, Zsolt Csabai, Zsolt Balázs, Emese Kis, Judit Molnár, and Zsolt Boldogkői. Third-generation sequencing reveals extensive polycistronism and transcriptional overlapping in a baculovirus. Sci Rep, 8(1): 8604, 2018.

[36] Rachael E Workman, Alison Tang, Paul S Tang, Miten Jain, John R Tyson, Philip C Zuzarte, Timothy Gilpatrick, Roham Razaghi, Joshua Quick, Norah Sadowski, Nadine Holmes, Jaqueline Goes de Jesus, Karen Jones, Terrance P Snutch, Nicholas James Loman, Benedict Paten, Matthew W Loose, Jared T Simpson, Hugh E Olsen, Angela N Brooks, Mark Akeson, and Winston Timp. Nanopore native RNA sequencing of a human poly(A) transcriptome. bioRxiv, page 459529, 2018.

[37] Daniel P Depledge, Srinivas Kalanghad Puthankalam, Tomohiko Sadaoka, Devin Beady, Yasuko Mori, Dimitris Placantonakis, Ian Mohr, and Angus Wilson. Native RNA sequencing on nanopore arrays redefines the transcriptional complexity of a viral pathogen. bioRxiv, page 373522, 2018.

[38] Matthew W Keller, Benjamin L Rambo-Martin, Malania M Wilson, Callie A Ridenour, Samuel S Shepard, Thomas J Stark, Elizabeth B Neuhaus, Vivien G Dugan, David E Wentworth, and John R Barnes. Direct RNA sequencing of the complete influenza A virus genome. bioRxiv, page 300384, 2018.

[39] Marcus H Stoiber, Joshua Quick, Rob Egan, Ji Eun Lee, Susan E Celniker, Robert Neely, Nicholas Loman, Len Pennacchio, and James B Brown. De novo identification of DNA modifications enabled by genome-guided nanopore signal processing. bioRxiv, page 094672, 2016.

[40] Alexa BR McIntyre, Noah Alexander, Aaron S Burton, Sarah Castro-Wallace, Charles Y Chiu, Kristen K John, Sarah E Stahl, Sheng Li, and Christopher E Mason. Nanopore detection of bacterial DNA base modifications. bioRxiv, page 127100, 2017.

[41] Ramakanth Madhugiri, Nadja Karl, Daniel Petersen, Kevin Lamkiewicz, Markus Fricke, Ulrike Wend, Robina Scheuer, Manja Marz, and John Ziebuhr. Structural and functional conservation of cis-acting RNA elements in coronavirus 5’-terminal genome regions. Virology, 517:44–55, 2018. ISSN 1096-0341. doi: 10.1016/j.virol.2017.11.025.

[42] S F Altschul, W Gish, W Miller, E W Myers, and D J. Lipman. Basic local alignment search tool. J Mol Biol, 215:403–10, 1990.

[43] Alexander O Pasternak, Willy J M Spaan, and Eric J Snijder. Nidovirus transcription: how to make sense…? J Gen Virol, 87: 1403–1421, 2006. ISSN 0022-1317. doi: 10.1099/vir.0.81611-0.

[44] Isabel Sola, Fernando Almazan, Sonia Zuniga, and Luis Enjuanes. Continuous and discontinuous RNA dynthesis in coronaviruses. Annu Rev Virol, 2:265–288, 2015. ISSN 2327-0578. doi: 10.1146/annurev-virology-100114-055218.

[45] W Luytjes, H Gerritsma, and W J Spaan. Replication of synthetic defective interfering rnas derived from coronavirus mouse hepatitis virus-A59. Virology, 216:174–183, 1996. ISSN 0042-6822. doi: 10.1006/viro.1996.0044.

[46] Karel Břinda, Valentina Boeva, and Gregory Kucherov. Ococo: an online consensus caller. arXiv, 2017.

[47] Sergey Koren, Brian P. Walenz, Konstantin Berlin, Jason R. Miller, Nicholas H. Bergman, and Adam M. Phillippy. Canu: scalable and accurate long-read assembly via adaptive *k* -mer weighting and repeat separation. Genome Res, 27(5):722–736, mar 2017. doi: 10.1101/gr.215087.116.

[48] Pierre Morisse, Thierry Lecroq, and Arnaud Lefebvre. Hybrid correction of highly noisy long reads using a variable-order de Bruijn graph. Bioinformatics, 2018. ISSN 1367-4811. doi: 10.1093/bioinformatics/bty521.

[49] David A Brian and Willy JM Spaan. Recombination and coronavirus defective interfering RNAs. Semin Virol, 8(2):101–111, 1997.

[50] Z Penzes, K Tibbles, K Shaw, P Britton, T D Brown, and D Cavanagh. Characterization of a replicating and packaged defective RNA of avian coronavirus infectious bronchitis virus. Virology, 203:286–293, 1994. ISSN 0042-6822. doi: 10.1006/viro.1994.1486.

[51] Z Penzes, C Wroe, T D Brown, P Britton, and D Cavanagh. Replication and packaging of coronavirus infectious bronchitis virus defective RNAs lacking a long open reading frame. J Virol, 70:8660–8668, 1996. ISSN 0022-538X.

[52] A Mendez, C Smerdou, A Izeta, F Gebauer, and L Enjuanes. Molecular characterization of transmissible gastroenteritis coronavirus defective interfering genomes: packaging and heterogeneity. Virology, 217:495–507, 1996. ISSN 0042-6822. doi: 10.1006/viro.1996.0144.

[53] A Izeta, C Smerdou, S Alonso, Z Penzes, A Mendez, J Plana-Duran, and L Enjuanes. Replication and packaging of transmissible gastroenteritis coronavirus-derived synthetic minigenomes. J Virol, 73:1535–1545, 1999. ISSN 0022-538X.

[54] Isabel Sola, Pedro A Mateos-Gomez, Fernando Almazan, Sonia Zuniga, and Luis Enjuanes. RNA-RNA and RNA-protein interactions in coronavirus replication and transcription. RNA Biol, 8: 237–248, 2011. ISSN 1555-8584.

[55] Hung-Yi Wu and David A Brian. Subgenomic messenger RNA amplification in coronaviruses. Proc Natl Acad Sci USA, 107: 12257–12262, 2010. ISSN 1091-6490. doi: 10.1073/pnas.1000378107.

[56] Sara Alonso, Ander Izeta, Isabel Sola, and Luis Enjuanes. Transcription regulatory sequences and mRNA expression levels in the coronavirus transmissible gastroenteritis virus. J Virol, 76: 1293–1308, 2002. ISSN 0022-538X.

[57] A Ozdarendeli, S Ku, S Rochat, G D Williams, S D Senanayake, and D A Brian. Downstream sequences influence the choice between a naturally occurring noncanonical and closely positioned upstream canonical heptameric fusion motif during bovine coronavirus subgenomic mRNA synthesis. J Virol, 75:7362–7374, 2001. ISSN 0022-538X. doi: 10.1128/JVI.75.16.7362-7374.2001.

[58] M M Lai and D Cavanagh. The molecular biology of coronaviruses. Adv Virus Res, 48:1–100, 1997. ISSN 0065-3527.

[59] M M Lai. Cellular factors in the transcription and replication of viral RNA genomes: a parallel to DNA-dependent RNA transcription. Virology, 244:1–12, 1998. ISSN 0042-6822. doi: 10.1006/viro.1998.9098.

[60] M Joo and S Makino. Mutagenic analysis of the coronavirus intergenic consensus sequence. J Virol, 66:6330–6337, 1992. ISSN 0022-538X.

[61] Achim Breiling and Frank Lyko. Epigenetic regulatory functions of DNA modifications: 5-methylcytosine and beyond. Epigenetics Chromatin, 8:24, 2015. ISSN 1756-8935. doi: 10.1186/s13072-015-0016-6.

[62] Ian A Roundtree, Molly E Evans, Tao Pan, and Chuan He. Dynamic RNA modifications in gene expression regulation. Cell, 169:1187–1200, June 2017. ISSN 1097-4172. doi: 10.1016/j.cell.2017.05.045.

[63] Anya V Grozhik and Samie R Jaffrey. Epitranscriptomics: Shrinking maps of RNA modifications. Nature, 551:174–176, November 2017. ISSN 1476-4687. doi: 10.1038/nature24156.

[64] Magdalena A Machnicka, Kaja Milanowska, Okan Osman Oglou, Elzbieta Purta, Malgorzata Kurkowska, Anna Olchowik, Witold Januszewski, Sebastian Kalinowski, Stanislaw Dunin-Horkawicz, Kristian M Rother, Mark Helm, Janusz M Bujnicki, and Henri Grosjean. MODOMICS: a database of RNA modification pathways–2013 update. Nucleic Acids Res, 41:D262–D267, January 2013. ISSN 1362-4962. doi: 10.1093/nar/gks1007.

[65] Nicholas J Loman, Joshua Quick, and Jared T Simpson. A complete bacterial genome assembled de novo using only nanopore sequencing data. Nat Methods, 12:733–735, 2015. ISSN 1548-7105. doi: 10.1038/nmeth.3444.

[66] Tobias Hertzig, Elke Scandella, Barbara Schelle, John Ziebuhr, Stuart G Siddell, Burkhard Ludewig, and Volker Thiel. Rapid identification of coronavirus replicase inhibitors using a selectable replicon RNA. J Gen Virol, 85(6):1717–1725, 2004.

[67] V Thiel, J Herold, B Schelle, and S G Siddell. Infectious RNA transcribed in vitro from a cDNA copy of the human coronavirus genome cloned in vaccinia virus. The Journal of general virology, 82:1273–1281, June 2001. ISSN 0022-1317. doi: 10.1099/0022-1317-82-6-1273.

[68] V Thiel and S G Siddell. Reverse genetics of coronaviruses using vaccinia virus vectors. Curr Top Microbiol Immunol, 287:199–227, 2005. ISSN 0070-217X.

[69] Heng Li. Minimap2: pairwise alignment for nucleotide sequences. Bioinformatics, 2018. ISSN 1367-4811. doi: 10.1093/bioinformatics/bty191.

[70] Stacia R Engel, Fred S Dietrich, Dianna G Fisk, Gail Binkley, Rama Balakrishnan, Maria C Costanzo, Selina S Dwight, Benjamin C Hitz, Kalpana Karra, Robert S Nash, Shuai Weng, Edith D Wong, Paul Lloyd, Marek S Skrzypek, Stuart R Miyasato, Matt Simison, and J Michael Cherry. The reference genome sequence of *Saccharomyces cerevisiae*: then and now. G3 (Bethesda), 4: 389–398, March 2014. ISSN 2160-1836. doi: 10.1534/g3.113.008995.

