## Supplemental Figure S1 for "Direct RNA nanopore sequencing of full-length coronavirus genomes provides novel insights into structural variants and enables modification analysis"

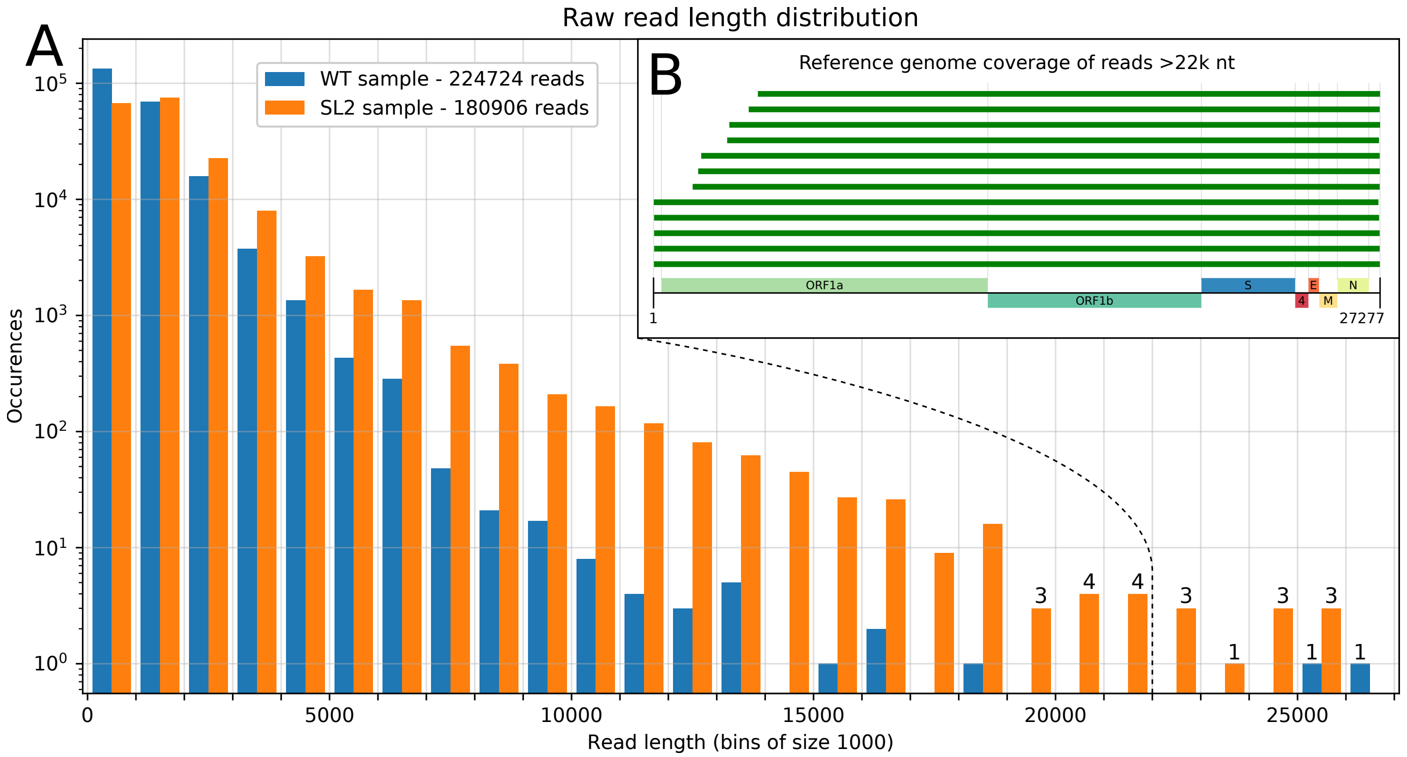


**Supplemental Figure 1** **A:** Length distribution histogram of basecalled reads in both samples. The WT sample produced more reads overall, but the majority of reads was shorter than 1,000 nt (median read length: 826). In the SL2 sample, the sequenced reads were generally longer (median read length: 1342). The difference was most pronounced for very long reads, in particular for lengths greater than 10k nt. In total, 5,543 reads were longer than 5k nt and 594 reads were longer than 10k nt. **B:** Spanned regions of the reference genome for the 12 longest reads (>22k nt) from both data sets. Five of these cover almost the complete genome and thus represent full viral RNA genomes, sequenced directly without a cDNA step.
