## Supplemental Figure S2 for "Direct RNA nanopore sequencing of full-length coronavirus genomes provides novel insights into structural variants and enables modification analysis"

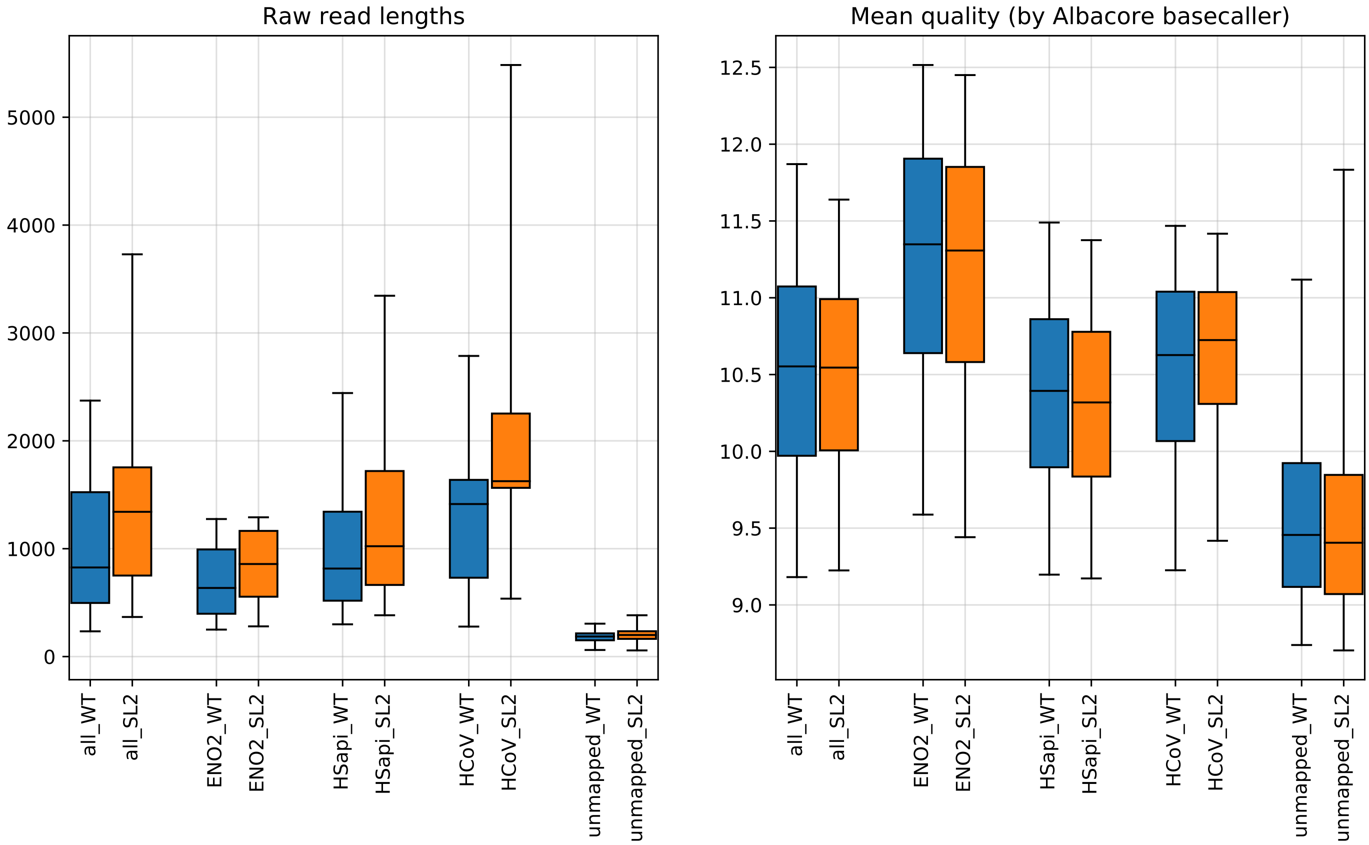


**Supplemental Figure 1** Box plots for raw properties of different read groups (mapping target categories based on minimap2 alignments). Boxes show quartiles, whiskers show 5/95 percentiles. Note that extreme values are omitted for visibility reasons (the very long reads covering the full virus genome would dominate the y-axis). **Left:** Raw read lengths per category. Reads that were not aligned by minimap ('unmapped') are mostly very short (median length <200). **Right:** Mean read quality by category. Reads in the 'unmapped' group display reduced quality. Note that the quality score given to each base by the Albacore basecaller is not a Phred quality score for sequencing, but rather represents a confidence score of that particular basecall. Also note that reads aligned to the yeast enolase 2 mRNA show elevated average quality, which is expected because the `Albacore` basecaller was trained on this calibration strand. Due to the outlined quality concerns for reads in the 'unmapped' category, and because these represented only 0.30 % and 0.07 % of total bases respectively, we chose to only use the higher quality reads that minimap2 could align.
