## Supplemental Figure S3 for "Direct RNA nanopore sequencing of full-length coronavirus genomes provides novel insights into structural variants and enables modification analysis"

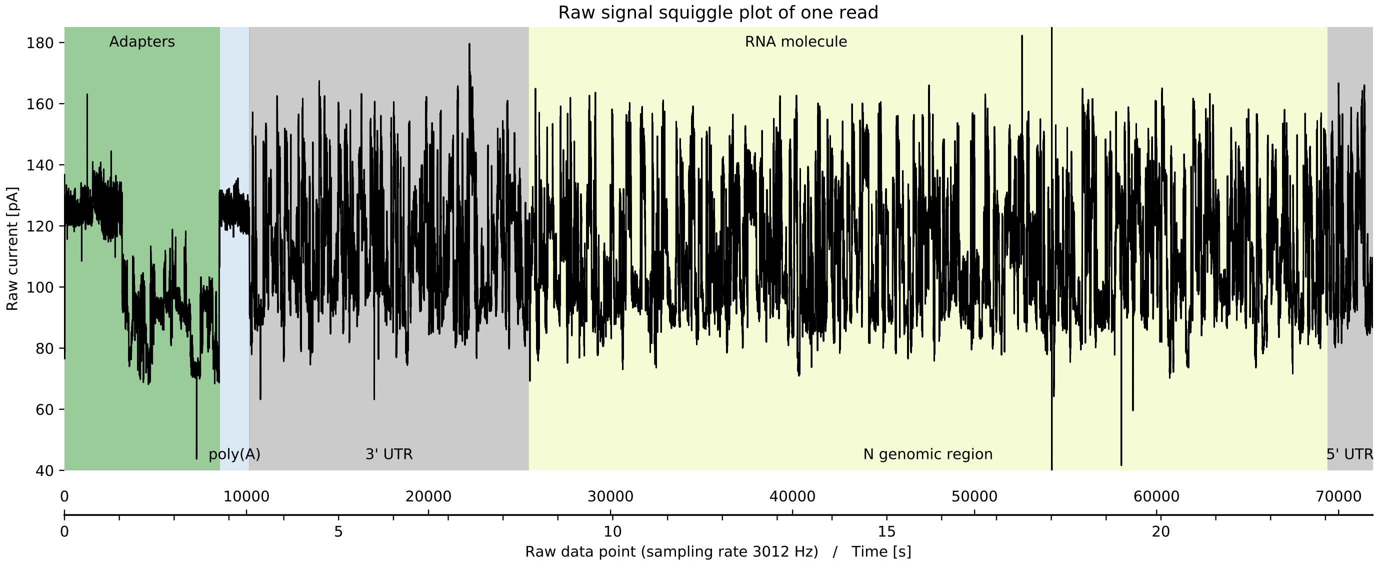


**Supplemental Figure 1** Raw signal 'squiggle' plot of one example read from the WT sample. Of note, sequencing starts from the 3'-polyadenylated end of the mRNA. The MinION measures the current inside the nanopore with a fixed sampling rate while a molecule passes through. Basecalling algorithms work directly on this raw signal data to infer the nucleotide sequence (Garalde *et al.* 2018). Unlike all other current sequencing technologies, nanopore sequencing conserves the information about base modifications in the raw signal (Garalde *et al.* 2018). SFig. 2 exemplarily shows a 5mC methylation pattern detected on raw signal data.

### References

Garalde, D. R., E. A. Snell, D. Jachimowicz, B. Sipos, J. H. Lloyd, *et al.*, 2018 Highly parallel direct RNA sequencing on an array of nanopores. Nat Methods **15**: 201–206.
