## Supplemental Figure S4 for "Direct RNA nanopore sequencing of full-length coronavirus genomes provides novel insights into structural variants and enables modification analysis"

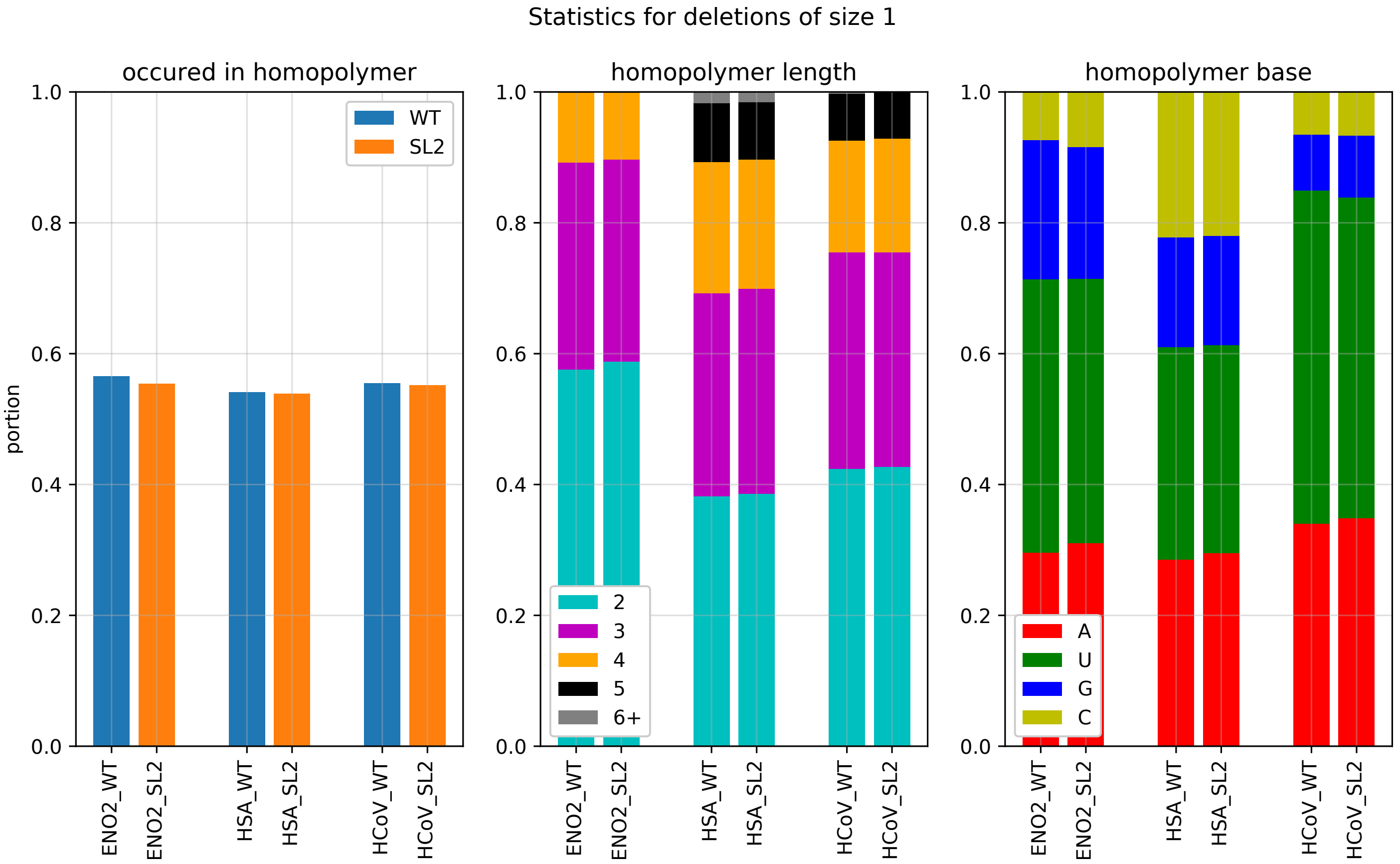


**Supplemental Figure 1** Statistics for all deletions of size 1 in all mappings against the yeast enolase 2, human and HCoV. **Left:** More than 53 % of all size 1 deletions occured in homopolymers (minimum stretch length of 2). **Middle:** A large proportion of the homopolymer stretches that coincide with a deletion are longer than 2 nucleotides. **Right:** A/U stretches appear more prone to causing deletion errors during basecalling, even with consideration of the G/C content of HCoV at 38.3 %.
