## Supplemental Figure S5 for "Direct RNA nanopore sequencing of full-length coronavirus genomes provides novel insights into structural variants and enables modification analysis"

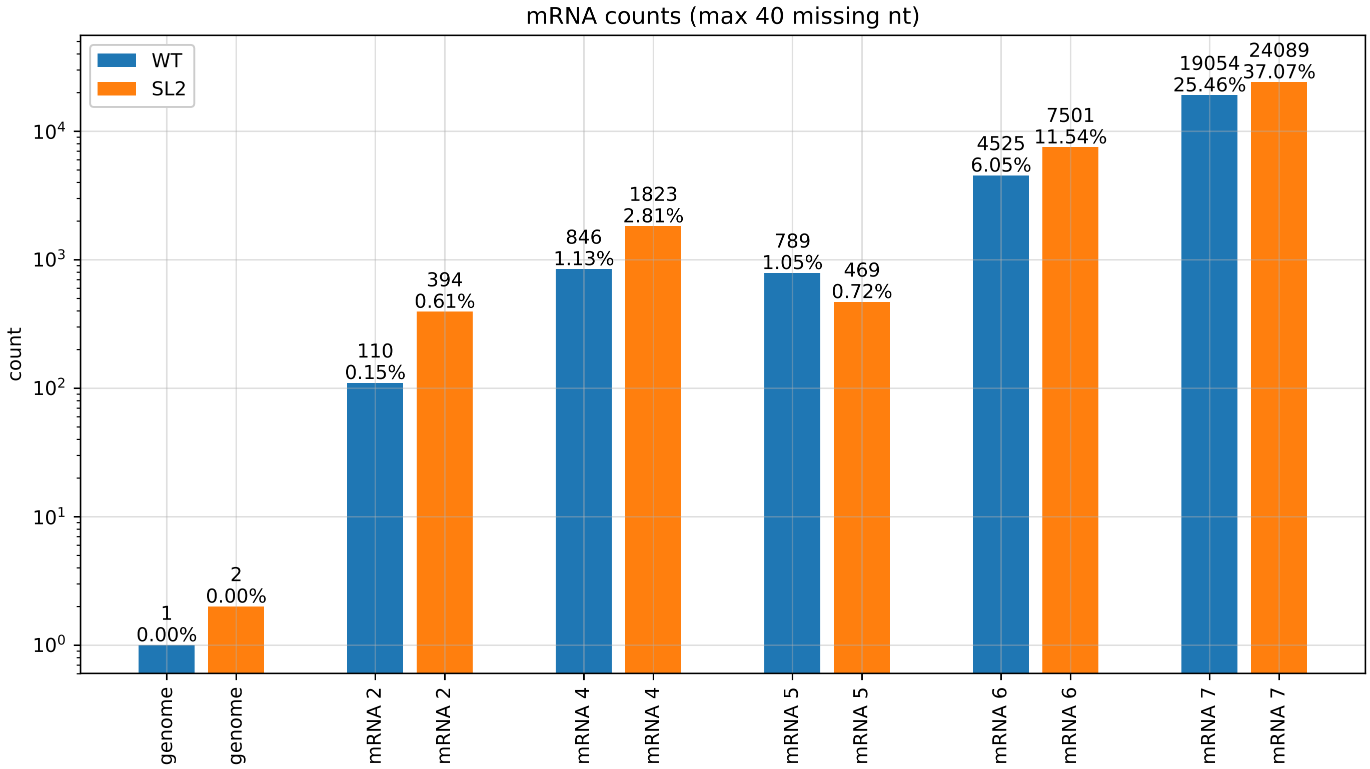


**Supplemental Figure 1** Number of mRNA copies in both samples, obtained by aligning all reads to the different mRNAs of HCoV (including the leader sequence, with exchanged stem loop 2 sequence for the SL2 sample). Aligned reads were considered copies of respective mRNAs when they were missing at most 40 nucleotides from both ends combined. Relative counts correspond accurately to the observed peaks in the alignment length distribution plot (Fig. 5). Note that 'genome' denotes the mRNA 1, i.e. the full genome.
