## Supplemental Figure S6 for "Direct RNA nanopore sequencing of full-length coronavirus genomes provides novel insights into structural variants and enables modification analysis"

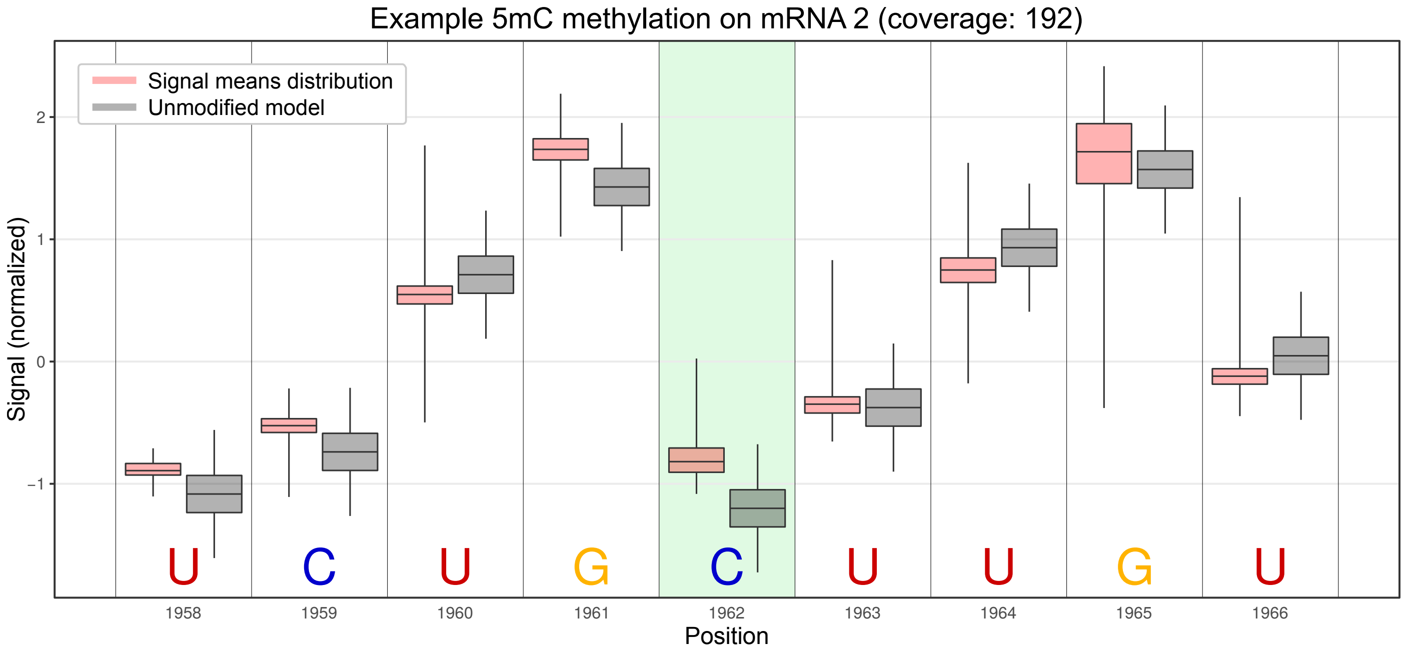


**Supplemental Figure 1** Example 5mC methylation pattern detected on raw signal data. The `Tombo` (Stoiber *et al.* 2016) resquiggle algorithm re-aligns the raw nanopore read signal to the basecalled read sequence. Subsequently, the raw signal at each nucleotide position (displayed as **red** distributions of event mean values) is tested with learned modification models (5mC for RNA) against baseline signal models representing unmodified bases (displayed as **grey** distributions). For the cytosin position highlighted in **green**, a methylation rate of 98.14 % was predicted across all 192 covering reads. Signals for surrounding bases are also affected by a modification, as the measured current is dependent on about 6 nucleotides that are located inside the nanopore at a time.

### References

Stoiber, M. H., J. Quick, R. Egan, J. E. Lee, S. E. Celniker, *et al.*, 2016 De novo identification of DNA modifications enabled by genome-guided nanopore signal processing. bioRxiv p. 094672.
