## Supplemental Figure S7 for "Direct RNA nanopore sequencing of full-length coronavirus genomes provides novel insights into structural variants and enables modification analysis"

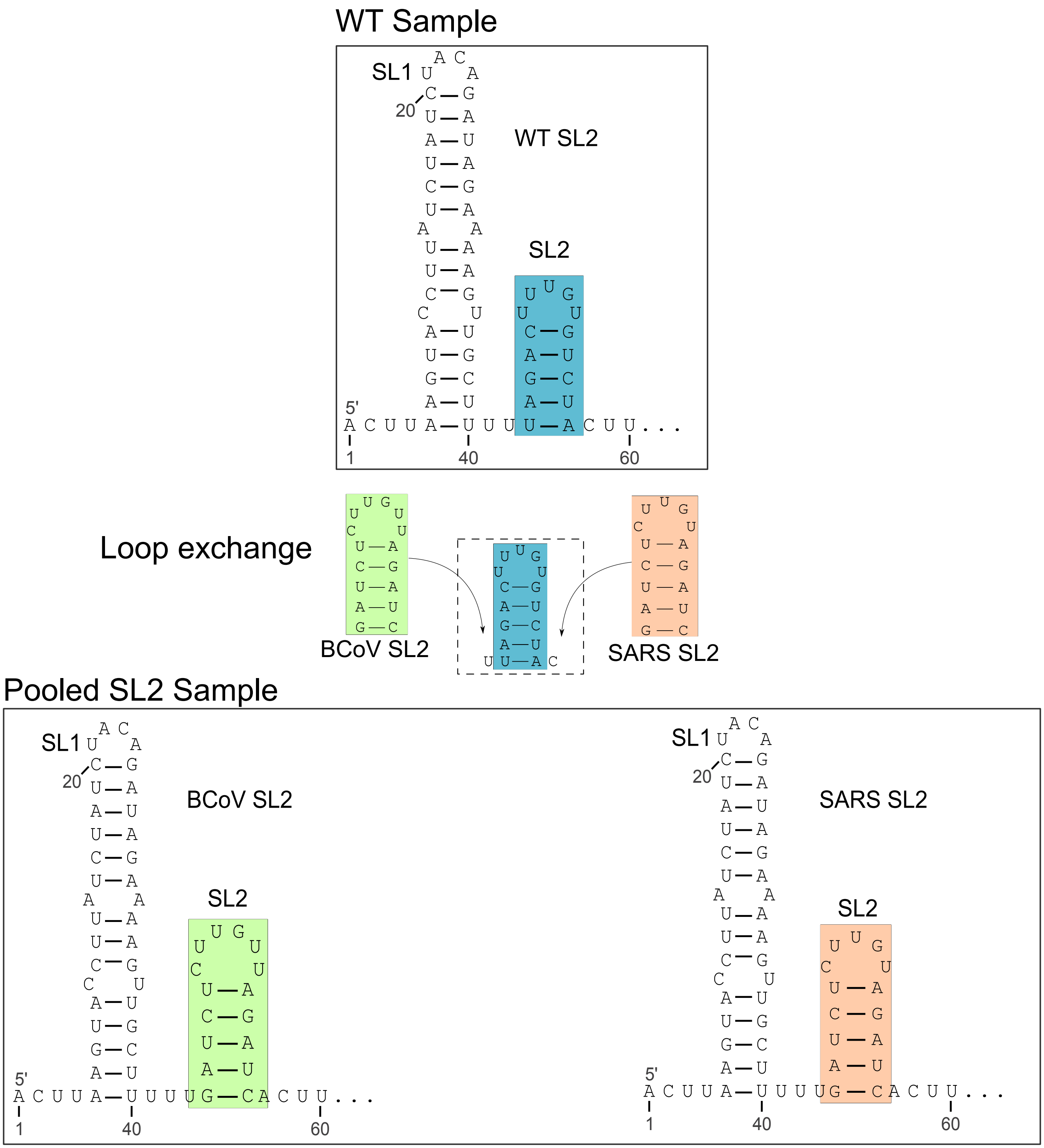


**Supplemental Figure 1 Upper:** The first 60 nucleotides of the 5′-UTR of the HCoV-229E are shown with their predicted RNA secondary structure using `RNAfold` (Lorenz *et al.* 2011). **Middle:** The SL2 region of the wild type (blue box) was exchanged with the BCoV (green box) and SARS-CoV (orange box) counterpart, respectively (Madhugiri *et al.* 2018). **Lower:** The resulting structure after changing the SL2 region.

### References

Lorenz, R., S. H. Bernhart, C. Höner Zu Siederdissen, H. Tafer, C. Flamm, *et al.*, 2011 ViennaRNA package 2.0. Algorithms Mol Biol **6**: 26.

Madhugiri, R., N. Karl, D. Petersen, K. Lamkiewicz, M. Fricke, *et al.*, 2018 Structural and functional conservation of cis-acting RNA elements in coronavirus 5′-terminal genome regions. Virology **517**: 44–55.
