## Supplemental Figure S8 for "Direct RNA nanopore sequencing of full-length coronavirus genomes provides novel insights into structural variants and enables modification analysis"

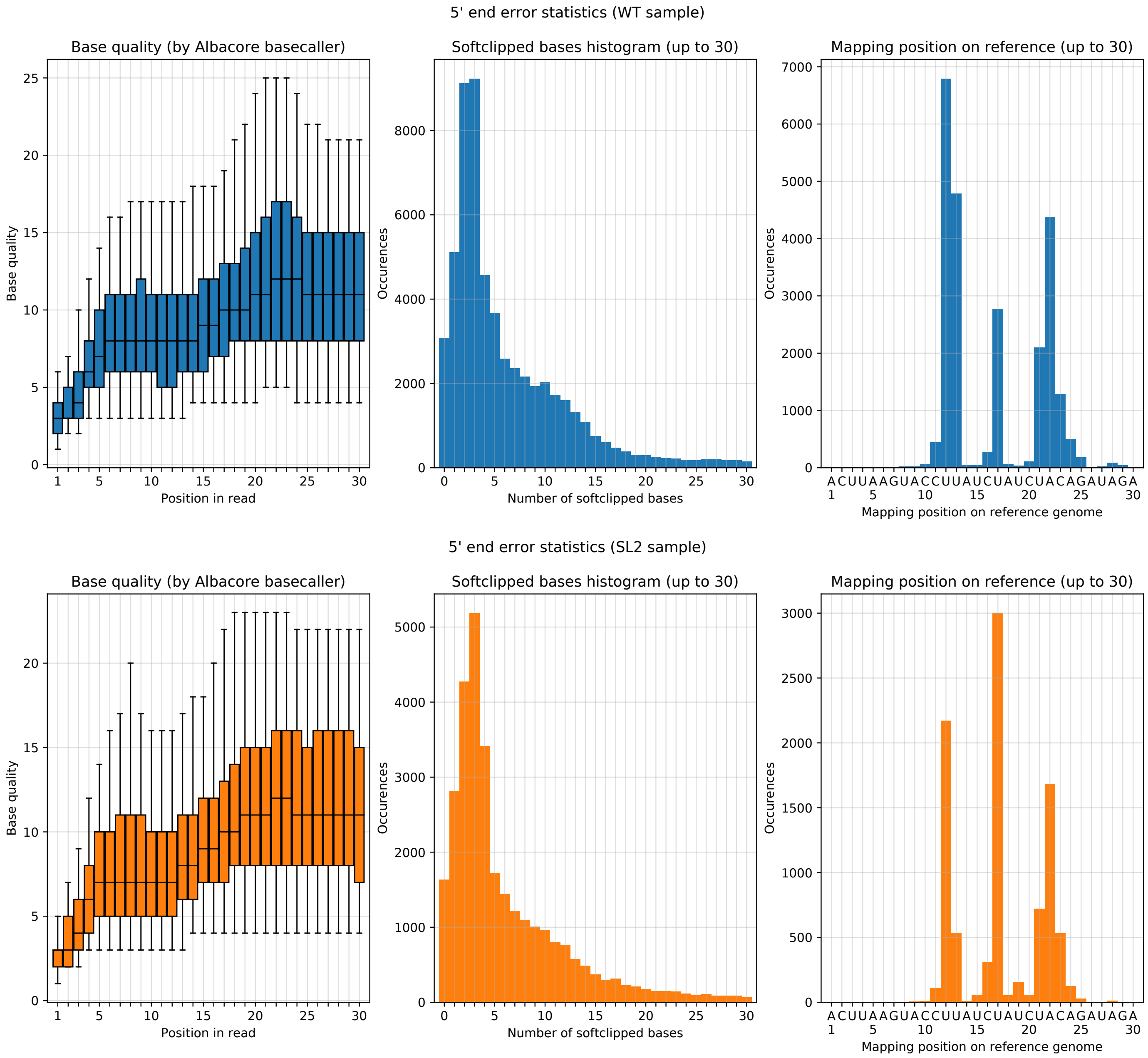


**Supplemental Figure 1** Statistics for the 5′ ends of reads. **Left:** As the RNA strands are sequenced in 3′ -> 5′ direction, the very last bases to pass the nanopore are those on the 5′ end. Basecalling these positions is plagued by reduced accuracy, probably due to less available context towards the end. In particular, the very last bases show poor accuracy, which is reflected in the quality score (basecalling confidence) given by the basecaller. While this effect is most extreme for only a few bases, a pattern of reduced confidence up to 20 bases long can be observed. **Middle:** As 5′ end basecalls are often false, almost all reads are not fully aligned on their 5′ end, and these bases are softclipped (i.e. skipped in alignment) by minimap2 during mapping. Looking at all alignments to the 5′ end of the reference genomes, one can observe that due to this effect the very first bases are not covered. **Right:** Mapping positions cluster around certain positions (12, 17 and 22), possibly due to bases/motifs that allow higher certainty for their respective basecall.
