## Supplemental Figure S9 for "Direct RNA nanopore sequencing of full-length coronavirus genomes provides novel insights into structural variants and enables modification analysis"

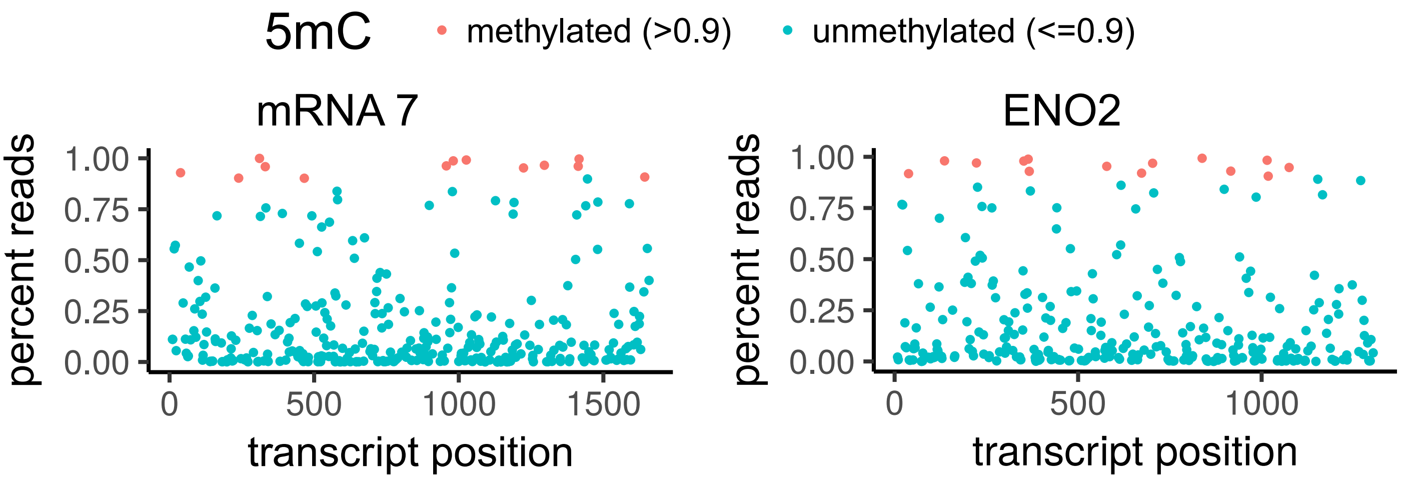


**Supplemental Figure 1** `Tombo` results for mRNA 7 and yeast ENO2. **Left:** Several cytosine positions were called as 5mC methylated in more than 90 % of covering reads. **Right:** The unmodified negative control – an *in vitro* yeast enolase II transcript – presented several positions that were assigned a methylated state by `Tombo`. We used this control to obtain a threshold for the minimum percentage of reads in the methylated state, so as to obtain a FPR of 5 %. As discussed in the manuscript, this threshold is 90 % of reads. While the overall methylation pattern looks similar, we nevertheless find consistent methylation across different subgenomic RNA "types", i.e. methylated positions of mRNA 2 are mirrored in mRNA 4 etc (see Fig. 6).
